# A CRISPR-based genetic screen in *Bacteroides thetaiotaomicron* reveals a small RNA modulator of bile susceptibility

**DOI:** 10.1101/2023.07.03.547467

**Authors:** Gianluca Prezza, Chunyu Liao, Sarah Reichardt, Chase L. Beisel, Alexander J. Westermann

**Affiliations:** Helmholtz Institute for RNA-based Infection Research (HIRI), Helmholtz Centre for Infection Research (HZI), Wu rzburg, D-97080, Germany; Medical Faculty, University of Wu rzburg, Wu rzburg, D-97080, Germany; Institute of Molecular Infection Biology (IMIB), University of Wu rzburg, Wu rzburg, D-97080, Germany

## Abstract

Microbiota-centric interventions are limited by our incomplete understanding of the gene functions of many of its constituent species. This applies in particular to small RNAs (sRNAs), which are emerging as important regulators in microbiota species, yet tend to be missed by traditional functional genomics approaches. Here, we establish CRISPR interference (CRISPRi) in the abundant microbiota member *Bacteroides thetaiotaomicron* for genome-wide sRNA screens. By assessing the abundance of different protospacer-adjacent motifs, we identify the *Prevotella bryantii* B14 Cas12a as a suitable nuclease for CRISPR screens in these bacteria and generate an inducible Cas12a expression system. Using a luciferase reporter strain, we infer guide design rules and use this knowledge to assemble a computational pipeline for automated gRNA design. By subjecting the resulting guide library to a phenotypic screen, we uncover the previously uncharacterized sRNA BatR to increase susceptibility to bile salts, likely through the regulation of genes involved in *Bacteroides* cell surface structure. Our study lays the groundwork for unlocking the genetic potential of these major human gut mutualists and, more generally, for discovering hidden functions of bacterial sRNAs.

## INTRODUCTION

The human intestinal microbiota is dominated by two bacterial phyla, the Firmicutes and the Bacteroidetes (1). Among the latter, obligate anaerobic gut *Bacteroides* are of paramount importance for human health and disease (2). To exploit these bacteria for the benefit of the host, we first need to decipher the functions of their genes. While we still have an incomplete understanding of the function of many protein-coding genes expressed by *Bacteroides* spp., there is barely any knowledge as to the function of noncoding genes, even in emerging microbiological model organisms such as *Bacteroides thetaiotaomicron* (3, 4).

The best understood and most prevalent class of noncoding RNA regulators in the bacterial kingdom are the small RNAs (sRNAs). These ∼50-250 nt-long RNA molecules typically regulate target gene expression by mediating base-pair interactions with complementary stretches within target mRNAs and control a wide range of cellular processes (5–7). For example, enterobacterial model species often deploy sRNAs to rapidly adapt their global gene expression to numerous intrinsic or environmental stress conditions (8). By contrast, only a handful of sRNAs have been characterized in *Bacteroides* spp. (9, 10). While functional genomics proved successful to link *Bacteroides* coding genes with phenotypes (11, 12), conventional approaches—such as transposon insertion-based perturbations (13, 14)—inactivate genes near-randomly. Consequently, in these approaches, the likelihood of a given gene being hit and inactivated is inversely proportional to its length. This renders such screens less practical for studying short genes, including genes encoding small proteins (15) or sRNAs. For example, of the currently annotated 135 intergenic *B*. *thetaiotaomicron* sRNAs, 54 were missed in a recent dense (315,668 unique mutants; on average, one insertion every ∼20 nucleotides) transposon insertion sequencing (TIS) screen (10, 12). By extrapolation, it follows that roughly 2.5 times more mutants would be minimally required for a transposon library to hit all annotated sRNAs in this organism. Even if such a library would be available, parallel screening of that many mutants would heavily increase the already large bottleneck effects that such approaches suffer from (14).

Instead, CRISPR interference (CRISPRi)—wherein specific guide RNAs (gRNAs) recruit catalytically inactive CRISPR-associated (Cas) nuclease molecules to interfere with target gene transcription (16)—lends itself for the knockdown of short genes. The minimal requirement for CRISPRi-based target gene suppression is the presence of a protospacer adjacent motif (PAM), which is usually composed of only a few nucleotides. The resulting targeting scope thus entails the entire genome, with the added advantage of tailored knockdowns of predefined gene sets. Recently, using a catalytically-dead *Streptococcus pyogenes* Cas9 (dCas9) and gRNA expression vectors, a proof-of-principle was provided for CRISPRi in *Bacteroides* spp. (17). However, as of now the technology has not been exploited for functional screening in these important gut mutualists. More generally, in spite of ongoing attempts (18, 19), to our knowledge CRISPRi has not previously been used for a systematic assessment of sRNA-associated phenotypes in any bacterial species.

Here, we deploy CRISPRi to knock down the full complement of known intergenic sRNAs of *B*. *thetaiotaomicron* type strain VPI-5482 for fitness screening. Of six Cas nucleases considered for CRISPRi, *Prevotella bryantii* B14 Cas12a (Pb2Cas12) recognizes a PAM that is the most represented within the 135 annotated intergenic *B*. *thetaiotaomicron* sRNAs. Combining an inducible dPb2Cas12a expression system with a luciferase reporter strain, we infer guide design rules for CRISPRi in *B*. *thetaiotaomicron* and employ this knowledge to generate a computational pipeline for automated library design. We use the resulting guide library to identify sRNAs whose repression affects *Bacteroides* resilience to bile stress. Through the resulting screen, we identify the previously uncharacterized sRNA BatR, which regulates genes involved in *Bacteroides* cell surface biosynthesis/assembly and confers enhanced susceptibility to bile salts. Altogether, the present guide library bears potential to systematically uncover phenotypes for *Bacteroides* sRNAs under a variety of experimental conditions. More generally, our work lays the ground for targeted gene knockdown in these abundant human microbiota members.

## RESULTS

### CRISPR nuclease selection and induced expression

We recently annotated a total of 366 noncoding genes in the genome of *B*. *thetaiotaomicron* type strain VPI-5482 (10). This list includes 156 *cis*-encoded antisense RNAs and 28 5’ UTR-derived and 24 3′ UTR-derived sRNAs. Targeted disruption of these noncoding RNA types may potentially affect the expression of the correspondingly overlapping protein-coding genes, thereby complicating the interpretation of possible phenotypes. Instead, we here focused on the 135 *B*. *thetaiotaomicron* sRNAs that do not overlap any other annotated gene, to which we refer to as “intergenic” sRNAs (10).

The low GC content of *B*. *thetaiotaomicron* intergenic sRNAs (35% (20)) likely results in an underrepresentation of the 5′-NGG-3′ PAM recognized by the most frequently used Cas9 nuclease from *Streptococcus pyogenes* (SpCas9). In order to maximize the number of potential targeting sites, we screened the intergenic sRNAs for the presence of PAMs of additional, characterized CRISPR effector nucleases, including 5′-NNAGAA-3′ (recognized by Sth1Cas9 (21)), 5′-TTV-3′ (FnCas12a, Pb2Cas12a, Lb6Cas12a (22)), 5′-NNNNGATT-3′ (NmCas9 (23)), 5′-TTTV-3′ (AsCas12a (24)), and 5′-NGRR-3′ (SaCas9 (25)) (Fig. 1A). The SpCas9 PAM had a moderate abundancy, with most sRNAs having a limited number of Cas9-PAMs. In contrast, the Cas12a-specific PAM 5′-TTV-3′ was the most frequent and present at least 6 times in each sRNA. Overall, this PAM remained the top hit when expanding the search space to the entire *B*. *thetaiotaomicron* genome sequence (Supplementary Fig. S1A), proposing Cas12a as a suitable nuclease for genome-wide *Bacteroides* screens in general terms.

**Figure 1:**
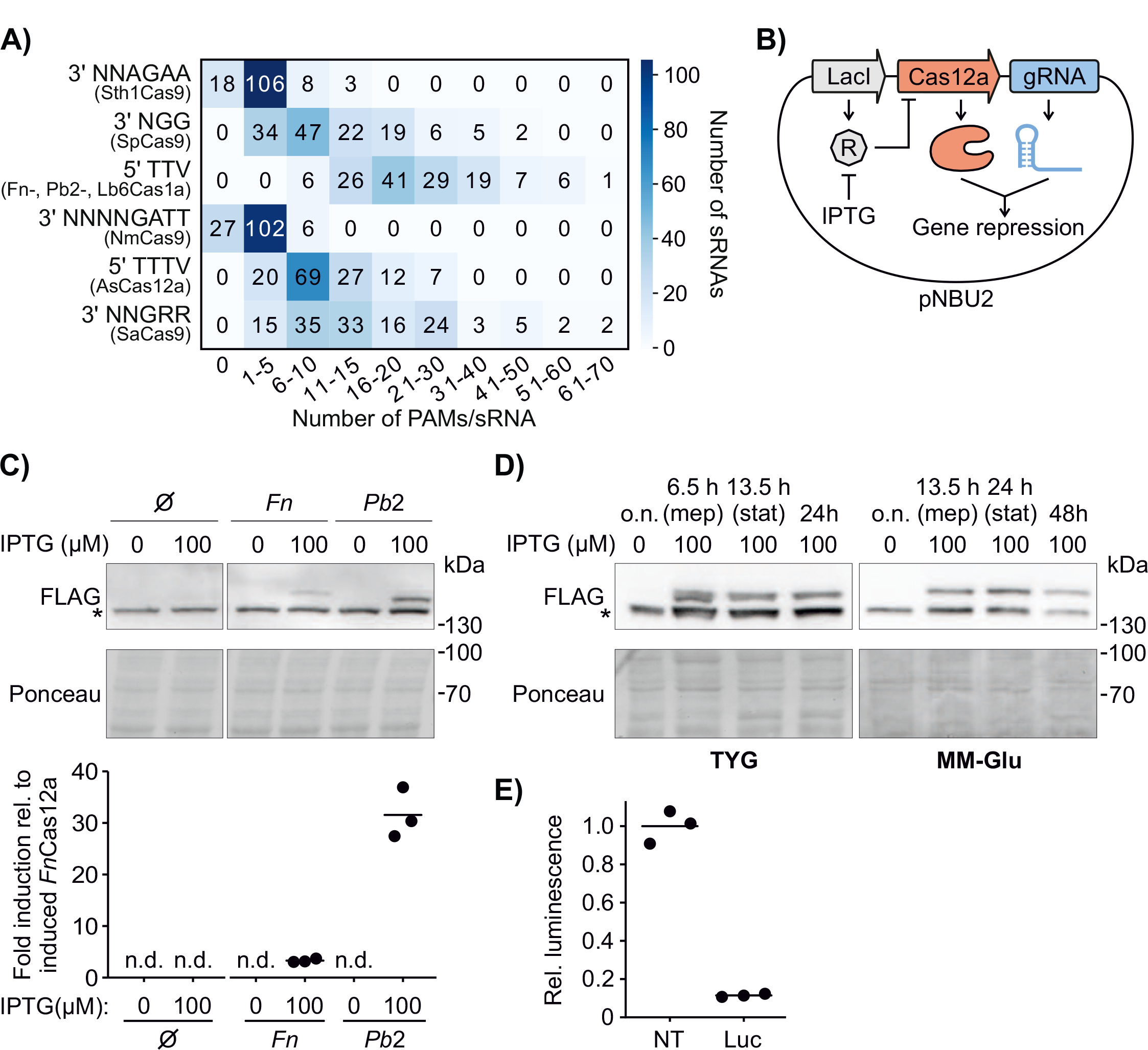
Nuclease selection for CRISPRi in *Bacteroides*. A) Frequency of PAM occurrences within the intergenic sRNAs of *B*. *thetaiotaomicron*. Color intensity is directly proportional to the number of sRNAs that contain the PAMs indicated on the y-axis. Examples of Cas nucleases recognizing each PAM are listed below the PAM sequence. B) Scheme of the construct for inducible and constitutive expression of Cas12a and gRNAs in *B*. *thetaiotaomicron*, respectively. The *lacI* gene produces a repressor (R) that blocks Cas12a expression. IPTG added to the culture media sequesters the repressor, allowing Cas12a transcription. A downstream gRNA is expressed from a constitutive promoter. Cas12a and the gRNA form an RNP complex that represses target gene transcription. C) Top: expression of FLAG-tagged FnCas12a and Pb2Cas12a in *B*. *thetaiotaomicron* at midexponential phase (OD_600_ 2, ∼6.5 h after IPTG induction). A sample harboring an empty construct (Ø) is included as a control. The asterisk next to the blot marks an unspecific band. Bottom: quantification of Cas12a band intensities (relative to induced FnCas12a). Horizontal lines represent the mean values of three biological replicates. n.d., not detected. D) Detection of Pb2Cas12a in *B*. *thetaiotaomicron* in rich (left, TYG) or minimal medium with glucose as sole carbon source (right, MM-Glu) after initial IPTG induction. The immunoblot is representative of three biological replicates and the asterisk marks an unspecific band. E) Luciferase activity after expression of a non-targeting control gRNA (NT) or a luciferasetargeting gRNA (Luc). Values are relative to the NT gRNA. Horizontal lines represent the means of three biological replicates.

Using an *E*. *coli* cell-free transcription-translation (TXTL) system (22, 26) as an initial screening tool, we tested the cleavage efficiency of three phylogenetically distinct Cas12a orthologs that recognize the same 5′-TTV-3′ PAM (*Lachnospiraceae bacterium* COE1, Lb6Cas12a; *Prevotella bryantii* B14, Pb2Cas12a; *Francisella novicida*, FnCas12a) (22) (Supplementary Fig. S1B). Of these orthologs, only Pb2Cas12a and FnCas12a yielded substantial cleavage of a targeted GFP reporter plasmid (Supplementary Fig. S1C). Next, we cloned Pb2Cas12a and FnCas12a into a pNBU2-based *Bacteroides* integration vector that allows expression of the nuclease from an IPTG-inducible promoter, while constitutively expressing a luciferase reporter gene and its corresponding gRNA (17) (Fig. 1B). After IPTG induction, both nucleases were detectable via western blotting, although Pb2Cas12a accumulated to a considerably higher level (Fig. 1C). We therefore selected this orthologue for CRISPRi. Upon induction with IPTG at a concentration of 250 µM—sufficient to achieve maximum expression of the nuclease (Supplementary Fig. S1D)—we detected the Pb2Cas12a protein at near-constant levels for at least 24 hours of growth in rich medium and for 48 hours in defined minimal medium with glucose (Fig. 1D).

CRISPRi uses a catalytically inactive nuclease variant to sterically interfere with RNA polymerase-mediated transcription without cleaving the targeted DNA. In analogy to the D917A variant that was previously found sufficient to inactivate the catalytic RuvC-like domain of FnCas12a (24), we introduced the D875A mutation into Pb2Cas12a (giving rise to ‘deactivated’ Pb2Cas12a [dPb2Cas12a]). Recruiting dPb2Cas12a to the promoter of the luciferase reporter construct resulted in a luminescence signal reduction of ∼10-fold (Fig. 1E). Together, these advances position dPb2Cas12a as a preferred nuclease for CRISPRi in *Bacteroides* for functional screening.

### Inferring gRNA design rules and construction of CRISPR arrays

As part of CRISPRi, the efficiency of gene knockdown is often dependent on the identity of the DNA strand that is targeted, with this specificity varying among Cas nucleases. Previous bacterial CRISPRi studies based on different Cas12a orthologs (FnCas12a, LbCas12a, AsCas12a) revealed that efficient knockdown can be achieved by targeting either strand within the promoter region (27–29). In contrast, when targeting within transcribed regions, knockdown was only effective when using gRNAs annealing to the template strand. To evaluate whether these findings can be extrapolated to the use of dPb2Cas12a in *B*. *thetaiotaomicron*, we designed eleven gRNAs complementary to either DNA strand within the promoter or coding region of the luciferase gene in our reporter strain (Fig. 2A). Compared to a non-targeting control gRNA, the gRNAs targeting either DNA strand in the promoter reduced the luciferase signal intensity by ∼3-10-fold. Within the coding region, gRNAs annealing to the template strand were more effective than those against the non-template strand. The only exception was gRNA ‘cds_NT1’, which anneals to the non-template strand in the 5′ portion of the coding region but still resulted in a 3-fold repression. This may be due to the proximity to the promoter region and/or the presence of the extended PAM ‘<u>T</u>TTV’, which is slightly preferred over ‘<u>V</u>TTV’ in *E*. *coli* (22). Based on these results, we concluded that in our system Cas12a-mediated transcriptional interference shows similar strand preferences than previously reported for other bacterial taxa and Cas12a orthologs.

**Figure 2:**
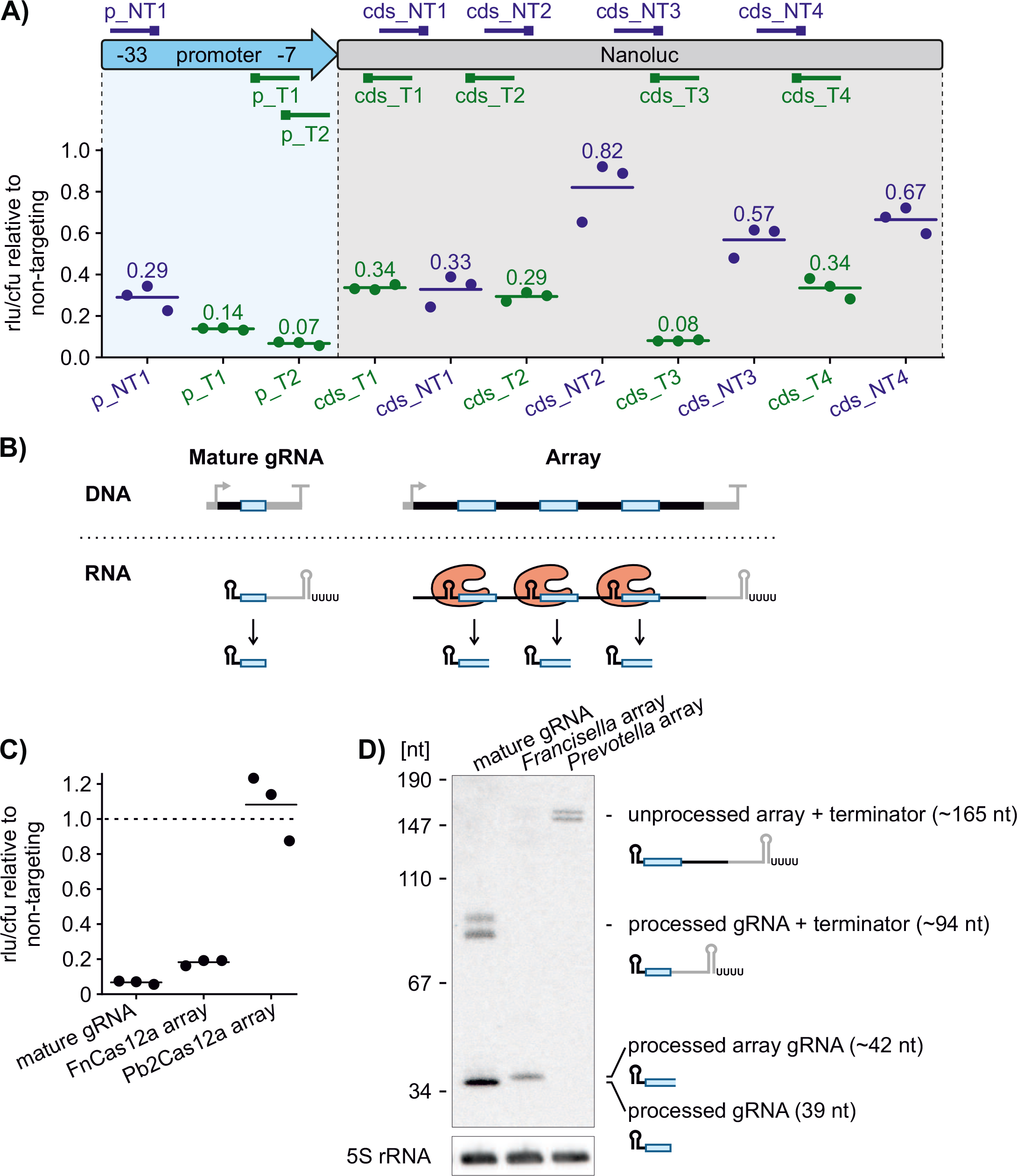
Targeting rules and array design for dPb2Cas12a. A) Knockdown of luciferase by dPb2Cas12a with different gRNAs. Top: scheme of the luciferase gene and promoter with relative position of the targeted regions. The position of the PAM is marked with a square. gRNAs shown in blue and green target the non-template and template strand, respectively. The -33 and -7 promoter regions are highlighted. Bottom: detected luciferase activity after expression of each gRNA. Values are relative to the levels observed in a strain coding a non-targeting gRNA. Horizontal lines mark the means of three biological replicates. B) Depiction of the expression of a single gRNA (left) or three gRNAs from an array (right). In the latter case, the single array transcript is recognized and processed by Cas12a into the three independent gRNA units. C) Knockdown of luciferase by dPb2Cas12a with the same spacer expressed as a gRNA (mature gRNA) or surrounded by repeats in a CRISPR array-like transcript (*Francisella* or *Prevotella* array). The data of the mature gRNA are the same shown in Fig. 2A for “p_T2”. Values are relative to levels observed in a strain with a non-targeting spacer, encoded either as gRNA, as an array with *Prevotella* repeats, or as an array with *Francisella* repeats (n=3). D) Detection by northern blotting of processed and precursor gRNA transcripts in the strains shown in panel C at mid-exponential phase (OD_600_ = 2.0, ∼6.5 h). The probe used is complementary to the spacer sequence. The *E*. *coli rnpB* terminator hairpin following the array/gRNA sequences gets post-transcriptionally removed (53, 54). The blot image is representative of three biological replicates.

One advantage of Cas12a over other Cas nucleases is its ability to process CRISPR arrays without the requirement of further accessory factors (30). Therefore, as an alternative to coexpressing mature gRNAs individually (Fig. 2B, left), expression of a CRISPR array would eventually result in the accumulation of multiple processed gRNAs starting from a single, compact transcript (Fig. 2B, right) (24). We explored to what extent this property of Cas12a can be exploited for simultaneous expression in *B*. *thetaiotaomicron* of multiple gRNAs derived from a single primary transcript. To this end, we designed an array composed of a spacer targeting the luciferase promoter (“p_T2” in Fig. 2A) surrounded by either *Francisella* or *Prevotella* Cas12a repeats, and we measured luminescence signals upon dPb2Cas12a induction. Surprisingly, only the array containing *Francisella* repeats—yet not the one with the *Prevotella* repeats—resulted in knockdown levels close to the ones achieved by delivering the mature gRNA (Fig. 2C). Northern blotting confirmed the accumulation of mature gRNAs when pro-cessed from the *Francisella* array (Fig. 2D). In contrast, no mature gRNAs were detected in cells encoding the *Prevotella* array (Fig. 2D), suggesting that *Prevotella* repeats are either inefficiently processed in *B*. *thetaiotaomicron* or improperly fold with the provided guide sequence to prevent Cas12a recognition (31). Taken together, dPb2Cas12a is generally capable of efficient gRNA maturation in *B*. *thetaiotaomicron*, albeit in a repeat-dependent manner. While the reason of the inefficient processing of *Prevotella* arrays remains elusive and our observations are only based on a single array, we nevertheless relied on *Francisella* repeats for all follow-on experiments.

### Efficient dCas12a-mediated inhibition of sRNA transcription and function

Building on the targeting rules inferred above, we designed gRNAs against two well-established *B*. *thetaiotaomicron* sRNAs, GibS (20) and MasB (10) (Fig. 3A, top). In each case, we designed three spacers annealing to different regions within the targeted sRNA gene or its promoter and expressed them as single gRNAs (‘g1–3′). When compared to a non-targeting control (‘gNT’), most of the gRNAs led to efficient (i.e., at least 3-fold) reduction in the respective sRNA levels (Fig. 3A, bottom), yet two gRNAs (‘GibS g2’ and ‘MasB g1’) resulted in negligible repression. While the cause of inefficient knockdown for these guides remains unknown, we note that by including multiple constructs targeting the same sRNA in the final mutant pool, the chances of failing to repress expression of a targeted sRNA are low.

**Figure 3:**
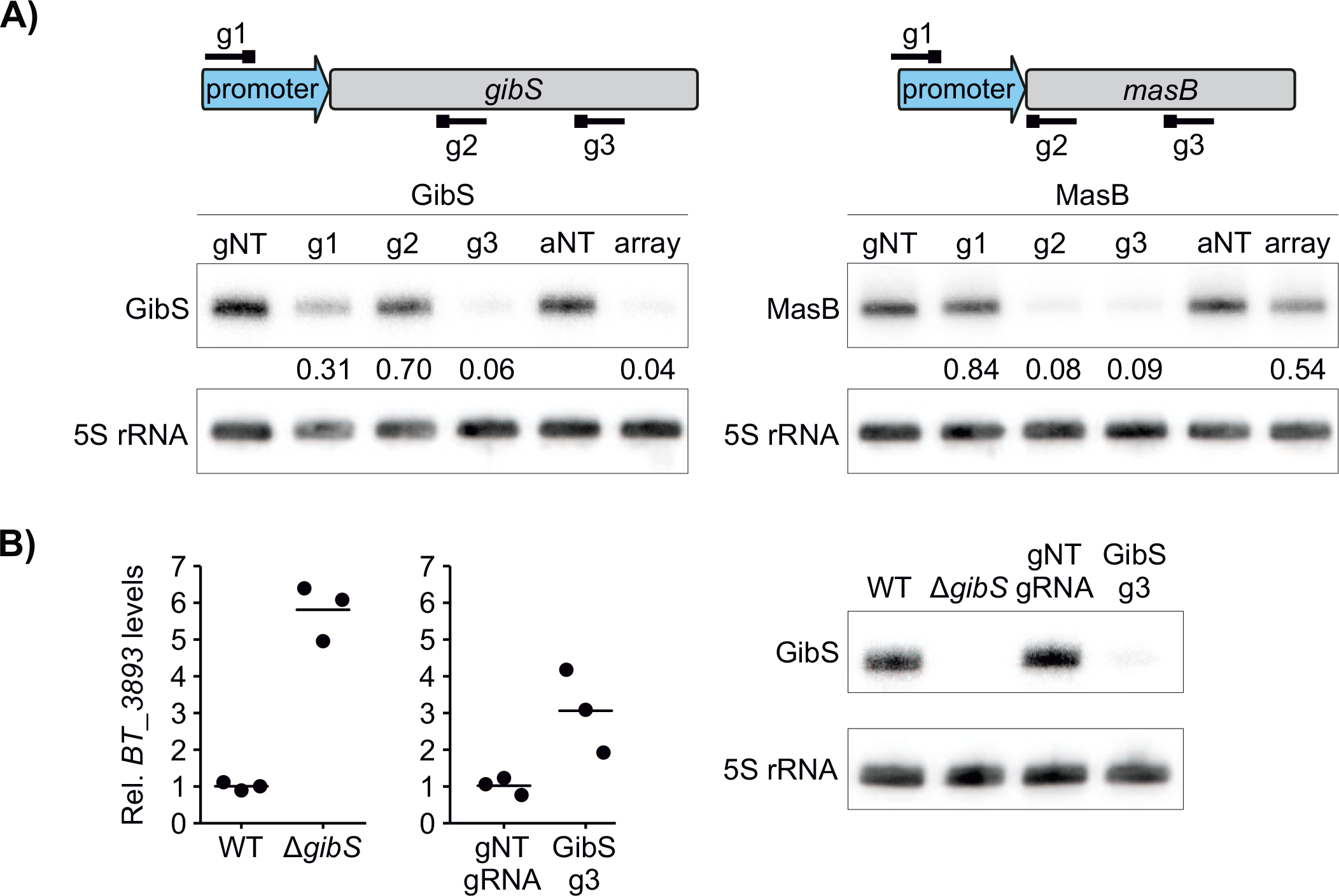
CRISPRi-mediated knockdown of selected *B*. *thetaiotaomicron* sRNAs. A) Depletion of the sRNAs GibS (left) and MasB (right) with dPb2Cas12a as detected by northern blotting. For each sRNA, three gRNAs (“g1-3”) or an array (“array”) were used, alongside a nontargeting gRNA (“gNT”) or non-targeting array (“aNT”). Relative annealing position of each spacer within the sRNA genes and promoters is depicted at the top. The blots shown are representative of three biological replicates and the mean value of each band intensity relative to the respective control is indicated below the blot. B) Recapitulation of a *gibS* deletion-associated phenotype with CRISPRi. Left: qRT-PCR measurements of *BT_3893* mRNA levels relative to the 16S rRNA in Δ*tdk* (WT), Δ*tdk*Δ*gibS* (Δ*gibS*) strains or in bacteria harboring the CRISPRi vector with a non-targeting (gNT gRNA) or a GibS-targeting (GibS g3) gRNA. Values shown are relative to Δ*tdk* or to the non-targeting gRNA-expressing strain, respectively. Horizontal lines represent the mean values from three biological replicates. Right: GibS levels as measured by northern blotting in the same samples. The blot is representative of three biological replicates.

We next assembled the three guides targeting the same sRNA into a CRISPR array. In the case of the *gibS*-targeting array (Fig. 3A, left), the steady-state level of the sRNA was reduced by ∼20-fold as compared to an array encoding a non-targeting guide (‘arrayNT’). However, the *masB*-targeting array failed to strongly suppress transcription of this sRNA (Fig. 3A, right), despite the fact that two of the contained guides led to a >10-fold repression when expressed as single gRNAs (‘g2, -3′). Processing of transcribed arrays by Cas12a involves the formation and recognition of a hairpin structure at the 3′ end of the repeats—a step essential for gRNA maturation (30). Prediction of the arrays’ secondary structure using *mfold* (32) revealed that all repeats in the *gibS*-targeting construct are predicted to correctly fold in at least one of the most probable (based on minimal free energy) structures. In contrast, in case of the *masB*-targeting array, several repeats did not form the terminal hairpin in any of the alternative structures (Supplementary Fig. S2A). Northern blotting with probes specific for each spacer of both arrays supported this prediction: all three mature anti-*gibS* gRNAs readily accumulated, while only one of the three anti-*masB* gRNAs was detected, and so at a very low level (Supplementary Fig. S2B). We concluded that, in line with previous observations (31), correct folding of the primary transcript must be taken into account when designing Cas12a arrays.

As a proof-of-concept for our CRISPRi-based sRNA knockdown approach in *B*. *thetaiotaomicron*, we tested whether expression of an anti-*gibS* gRNA would replicate molecular phenotypes previously established for a clean deletion mutant of this sRNA (20). Indeed, CRISPRi-mediated knockdown led to derepression of the confirmed GibS target *BT_3893* mRNA (Fig. 3B, left). We did not observe a similar relief in repression of another established GibS target, *BT_0771* (Supplementary Fig. S2C), in spite of reduced GibS levels (Fig. 3B, right). However, we hypothesize that *BT_0771* mRNA has a higher affinity to GibS than *BT_3893*, so residual traces of the sRNA that remain upon knockdown would be sufficient to repress *BT_0771* but not *BT_3893*. Collectively, these results show that sRNAs can be efficiently silenced in *B*. *thetaiotaomicron* via dPb2Cas12a-mediated CRISPRi, which can replicate phenotypes associated with clean sRNA deletions.

### Automated design of gRNAs against the full suite of intergenic B. thetaiotaomicron sRNAs

Based on the inferred design rules, we created a custom Python script to design a guide library against the 135 intergenic sRNAs of *B*. *thetaiotaomicron* (Fig. 4A). For each queried sRNA, the script designs three gRNAs that may anneal to either strand in the corresponding sRNA promoter (50 nts upstream of the transcription start site) or to the template strand within the transcribed region, yet discards all spacers that are prone to off-targeting (i.e., spacers with less than three mismatches to a second genomic locus and preceded by a 5′-TTV-3′ PAM). When multiple spacers are available, those with a preferred GC content (between 35 and 75%), annealing to the promoter or the 5′ portion of the targeted sRNA, or flanked by the preferred 5′-<u>T</u>TTV-3′ PAM (22) are prioritized. In addition, for each sRNA, our script attempts to design one array composed of three further, non-overlapping guides, taking correct array folding (see above) into account. In cases when array design is unsuccessful—either because of an insufficient number of available spacers, or due to improper folding of all possible arrays—only gRNAs are outputted. As a result, the script delivers a list of sequences: for gRNAs, spacers flanked by homology regions for Gibson assembly, and for each array, six sequences comprising both strands of each repeat-spacer unit flanked by compatible 4-nt overhangs for CRATES (31) assembly.

**Figure 4:**
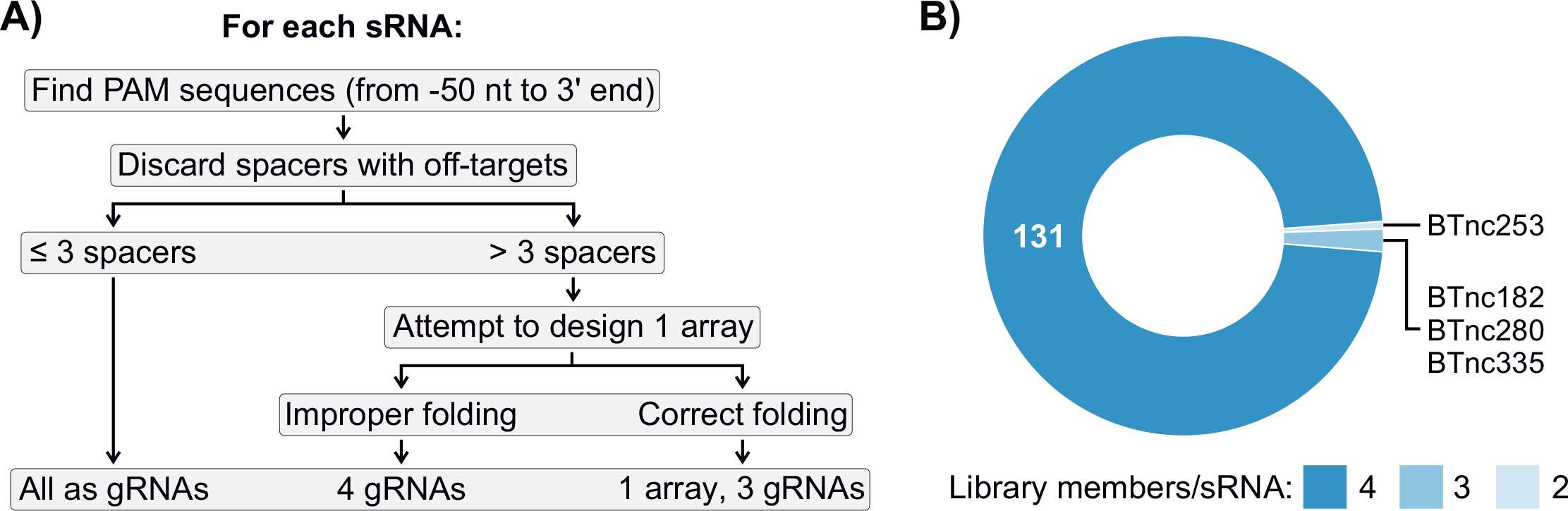
A computational pipeline for automated gRNA design and generation of a genomewide sRNA knockdown library. A) Flow chart of the script used to design the gRNAs/arrays targeting intergenic sRNAs of *B*. *thetaiotaomicron*. PAM sequences are identified for each sRNA in between 50 nt upstream of the transcription start site and the 3′ end of the transcript. Off-targeting spacers are discarded and gRNAs/arrays are designed. Arrays that are not predicted to form a hairpin at the 3′ end of each repeat are discarded. At the end the script outputs, when possible, four targeting constructs per sRNA. B) Representation of the sRNA knockdown mutants present in the final CRISPRi library.

Our script is freely available at https://github.com/gprezza/CRISPRi_tools and can be utilized to design libraries targeting any predefined gene set in *B*. *thetaiotaomicron* or any other species. Parameters such as PAM preference, strand-specificity, and array design are customizable, rendering the software suitable for versatile CRISPR applications. In the present context, a total of 577 gRNAs and 91 arrays were output by the script for the systematic knockdown of *Bacteroides* sRNAs. After cloning and conjugating the respective constructs into *B*. *thetaiotaomicron*, sequencing revealed that the vast majority (>99%) of the expected gRNAs and arrays were indeed contained in the resulting library. Most importantly, all 135 sRNAs were targeted in at least two library members, and only four sRNAs had fewer than the intended four gRNAs/arrays (Fig 4B; Supplementary Fig. S3). This guide library therefore lends itself to systematic sRNA mutant fitness screening.

### CRISPRi screening identifies BatR as a novel modulator of Bacteroides susceptibility to bile stress

Bile acids, whose amphipathic nature poses continuous stress to bacterial membranes (33), are important factors shaping the composition of the intestinal microbiota. As a test case for the utility of our CRISPRi approach for identifying biologically relevant noncoding RNAs, we screened for sRNAs that affect *Bacteroides* fitness in the presence of bile salts. To this end, we subcultured three independent uninduced overnight cultures of the input library into fresh TYG medium containing 250 μM of IPTG, either with or without 0.5 mg/mL of bile salts. Once the cultures reached stationary phase (∼13.5 h if unstressed and 24 h in the presence of bile salts (10)), we harvested the bacteria, extracted their genomic DNA, and PCR-amplified the full CRISPRi locus. After high-throughput sequencing, we compared the relative abundance of the guides in the presence or absence of bile stress, thereby deriving the relative fitness of each library member. To infer the effects of individual sRNAs, we averaged the fitness scores over different mutants targeting the same sRNA (Fig. 5A). This led to the prediction of two sRNAs whose silencing significantly affected *B*. *thetaiotaomicron* growth in the presence of bile salts (using an average fitness cutoff of |log_2_FC| > 0.5). Specifically, knockdown of BatR (formerly known as BTnc167 and here renamed to <u>B</u>ile <u>a</u>cid tolerance <u>R</u>egulator for reasons to follow) or BTnc353 resulted in increased or decreased bacterial numbers under bile stress, respectively, as compared with the bulk of the library members (Fig. 5B).

**Figure 5:**
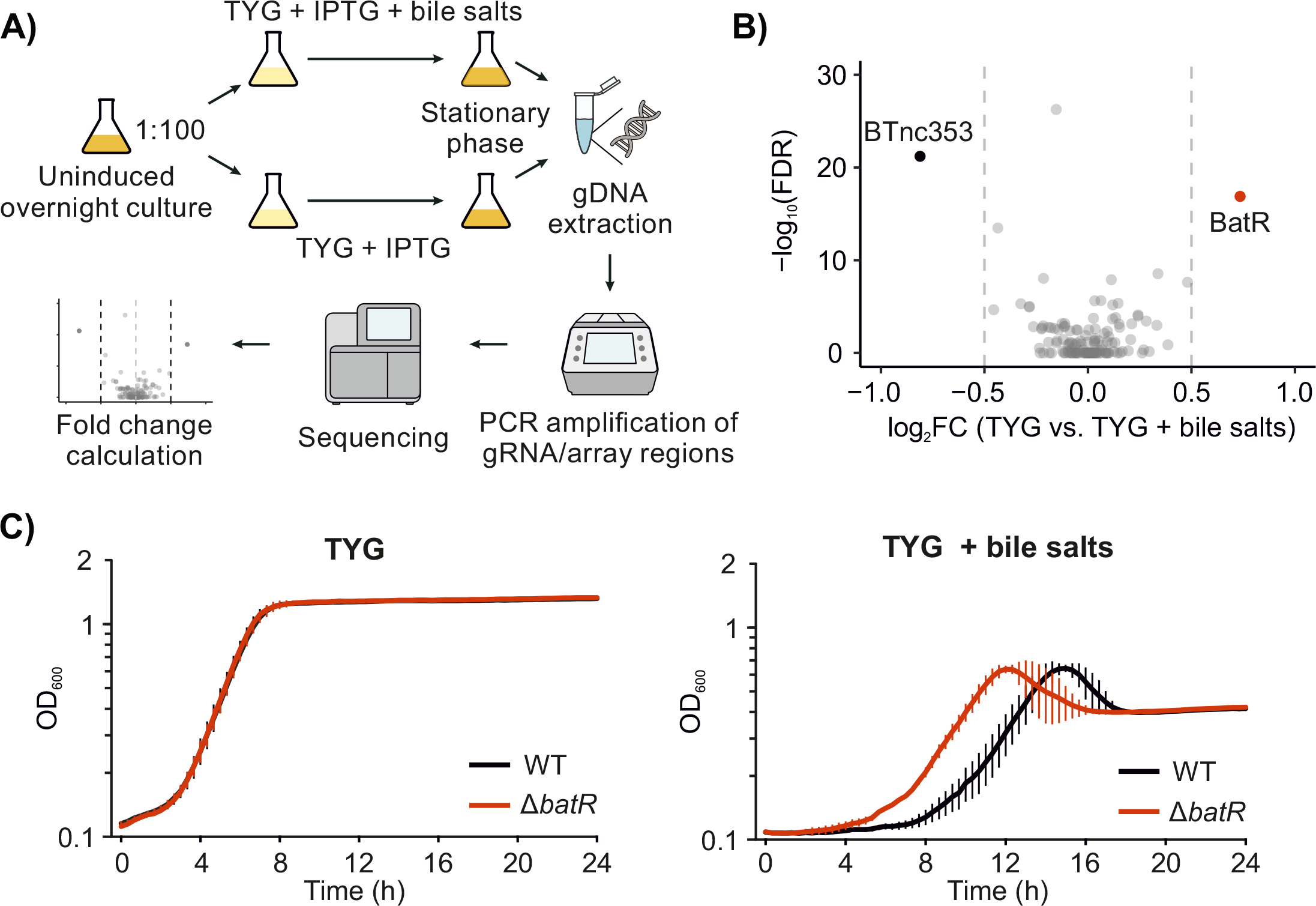
CRISPRi screening identifies an sRNA-associated bile stress phenotype. A) Workflow of the CRISPRi screen. An overnight culture of the library is diluted 1:100 into fresh medium with IPTG, with or without 0.05 mg/mL of bile salts. After selection, the region of the genome containing the gRNAs/arrays is PCR-amplified from both subcultures and sequenced, thus enabling quantification of the mutant strains and inferring a fitness score for each sRNA depletion under bile salts stress. B) Volcano plot of log_2_FC vs. false discovery rate of the per-sRNA summarized data from the bile salts selection experiment. sRNAs with log_2_FC > 0.5 and FDR < 0.05 are highlighted. Two dashed vertical lines mark the -0.5 and +0.5 log_2_FC thresholds. A positive log_2_FC means that mutants targeting the sRNA were on average enriched after growth in presence of bile salts, a negative log_2_FC indicates average depletion in the same condition. C) Growth of a clean Δ*batR* mutant and the wild-type strain in TYG medium without (left) or with (right) 0.05 mg/mL of bile salts. Error bars represent standard deviation of three biological replicates.

To validate the phenotype derived from the CRISPRi screen, we generated a clean deletion mutant of BatR and cultured it in TYG medium with or without 0.5 mg/mL of bile salts. As compared with an isogenic wild-type strain, the Δ*batR* strain did not display any growth difference in normal TYG, yet showed earlier entry into log phase when cultured in the presence of bile salts (Fig. 5C). This phenotype reflects the enhanced fitness observed in the CRISPRi screen, thus providing proof-of-principle for our approach to discover novel sRNA-associated fitness phenotypes.

### BatR is a conserved sRNA and involved in the regulation of cell surface genes

In light of the bile phenotype of the corresponding deletion mutant, we began to functionally characterize BatR. The sRNA itself and the corresponding –7 promoter box (20) are conserved in other *Bacteroides* species (Fig. 6A). Synteny analysis of the *batR* locus across *Bacteroides* spp. revealed its cooccurrence with uncharacterized transcriptional regulators and DNA-binding proteins (sigma factors, zinc finger proteins, Lrp/AsnC ligand binding domain-containing proteins), while a co-conservation with a set of isomerases, metallopeptidases, and methyltransferases was observed in most species other than *B*. *thetaiotaomicron* (Supplementary Fig. S4A). Assessment of our previously published transcriptomics data (10, 20) revealed that the sRNA is upregulated in exponentially growing bacteria, during carbon starvation, and after exposure to the secondary bile acid deoxycholate (Supplementary Fig. S4B). While the predicted length of BatR inferred from RNA-seq data was 374 nts (20), northern blotting revealed a transcript close to this length to be generally of low abundance in the cell (Fig. 6B, left). Instead, a 154 nt-long transcript and three shorter isoforms thereof accumulated, with the latter ones potentially deriving from 5′ processing events (‘p1,2,3′ processing sites, Fig. 6B, right). Based on these observations, we reannotated BatR in *B*. *thetaiotaomicron* as the 154 nt isoform. The predicted secondary structure of BatR comprises a strong Rho-independent terminator at the 3′ end, a second strong hairpin in central position, and a structured region near the 5′ end (Supplementary Fig. S4C, left). Processing at any of the three sites would likely render the region in between the two hairpins (position 86-109 in Supplementary Fig. S4C, left) single-stranded (Supplementary Fig. S4C, right) and hence, available for pairing with complementary stretches in target mRNAs.

**Figure 6:**
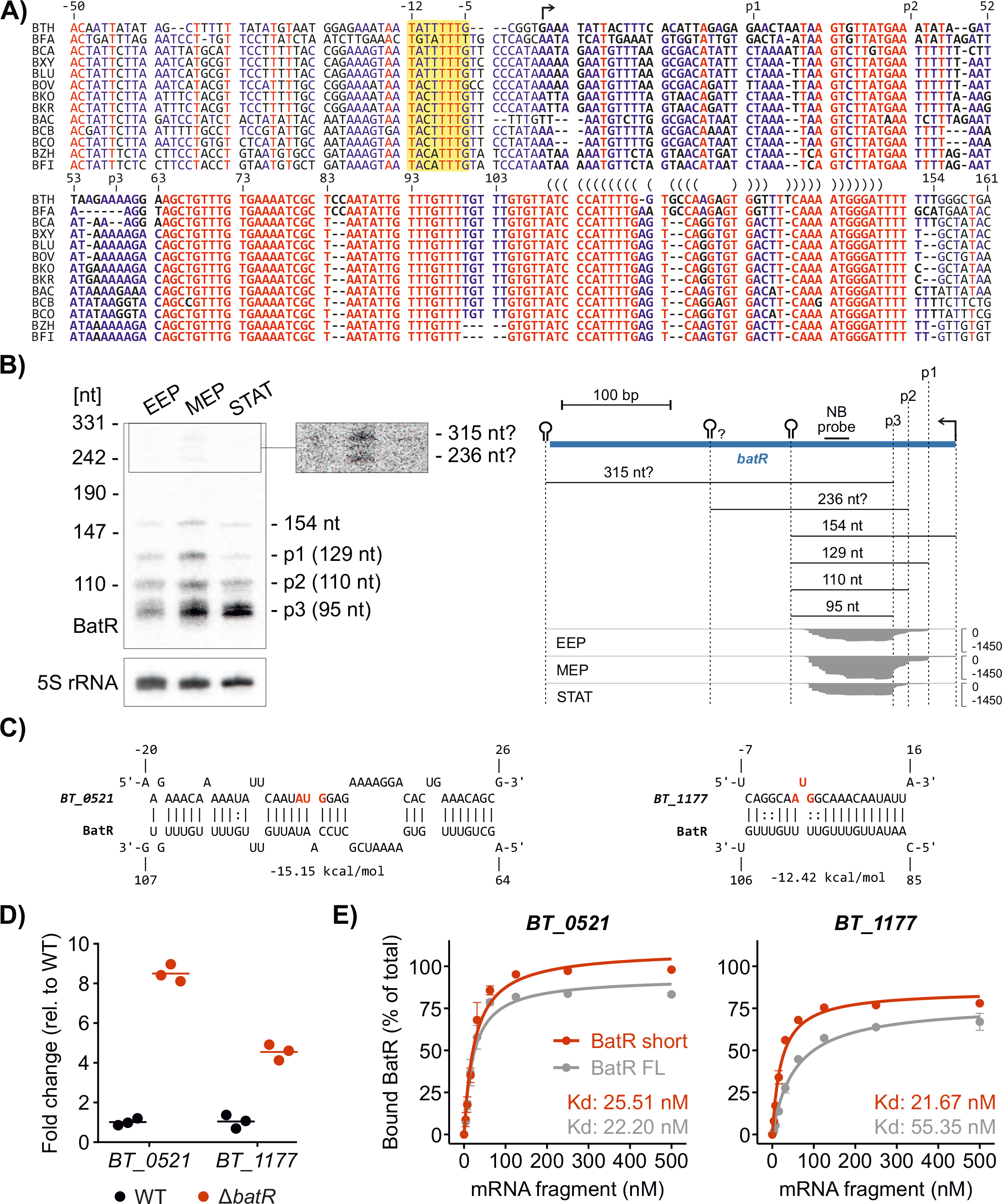
BatR characterization and target identification. A) Conservation of the *batR* genomic region spanning the four most prominent sRNA isoforms (see panel B). The -7 promoter box is highlighted in yellow, with the transcription start site indicated by an arrow. A bracket notation above the sequence marks the intrinsic terminator hairpin. BTH: *B*. *thetaiotaomicron*, BFA: *B*. *faecis*, BCA: *B*. *caecimuris*, BXY: *B*. *xylanisolvens*, BLU: *B*. *luhongzhouii*, BOV: *B*. *ovatus*, BKO: *B*. *koreensis*, BKR: *B*. *kribbi*, BAC: *B*. *acidifaciens*, BCB: *B*. *sp*. *CBA7301*, BCO: *B*. *congonensis*, BZH: *B*. *zhangwenhongii*, BFI: *B*. *finegoldii*. B) Left: northern blot-based validation of BatR size and expression profile in rich medium. Right: RNA-seq-derived expression profile of the annotated *batR* locus in rich medium. All isoforms observed on the northern blot are labeled and their sizes indicated. The previously annotated transcript contains two Rho-independent terminators that are marked with a hairpin symbol, and a possible third hairpin, additionally marked with a question mark. C) Interaction site between BatR and two putative mRNA targets as predicted by CopraRNA. The start codon is highlighted in red font. D) qRT-PCR measurement of the levels of selected BatR target candidates in the wild-type and Δ*batR* mutant. Horizontal lines represent the mean values of three biological replicates. E) Quantification of the fraction of bound, radiolabeled BatR at increasing concentrations of *BT_0521* (left) or *BT_1177* (right) from EMSAs shown in Supplementary Fig. S5B. Error bars represent standard deviation of two biological replicates. FL: full-length.

An *in*-*silico* target search using the CopraRNA tool (34) predicted that BatR binds the translation initiation region of a set of mRNAs encoding for proteins with a predicted role in cellsurface modulation (Fig. 6C, Supplementary Fig. S5A, Supplementary Table 5). Among the top ten target candidates (based on minimal free energy calculations upon BatR annealing), we picked four for further validation, namely *BT_0521* (partially homologous to the polysaccharide capsule synthesis protein CpsI from *Streptococcus iniae*), *BT_1177* (encoding an LPS biosynthesis protein), *BT_1195* (encoding a peptidoglycan transglycosylase), and *BT_3849* (encoding the LPS-assembly protein LptD). We collected total RNA from Δ*batR* bacteria or the isogenic wild-type cultured in TYG medium to midexponential phase (OD_600_ = 2). The ensuing qRT-PCR analysis revealed derepression of *BT_0521* (∼8fold) and *BT_1177* mRNA (∼5-fold), respectively, in the absence of BatR (Fig. 6D). Steady-state mRNA levels of the other two target candidates remained unchanged under the same conditions (Supplementary Fig. S5B). Electrophoretic mobility shift assays (EMSAs) using radiolabeled full-length (154 nt) BatR or its 95 nt isoform demonstrated direct binding to the 5′ regions of each of the above four predicted target candidates (Supplementary Fig. S5C). In line with our hypothesis that the sRNA’s seed region is partially occluded in full-length BatR but opens up upon processing, the dissociation constant (K*_d_*) and/or the fraction of bound sRNA at saturation were superior for the short isoform as compared with full-length BatR (Fig. 6E, Supplementary Fig. S5C). Taken altogether, our combined data suggest BatR as a novel post-transcriptional regulator of *Bacteroides* cell surface structure, with implications for the bacteria’s tolerance of bile stress, all based on genome-wide screening of sRNAs using an efficient CRISPRi system we established in *B*. *thetaiotaomicron*.

## DISCUSSION

Well-studied bacteria, such as the species belonging to the Enterobacteriaceae, employ sRNAs to cope with stresses and changing environmental conditions (8). Yet to what extent obligate anaerobic bacteria of the intestinal microbiota depend on sRNAs to thrive in the fluctuating niches of the gut remains an understudied branch of microbiome research. The Bacteroidetes represent a predominant phylum of human gut microbes (4). Recently, studies from us and others identified sRNAs in different gut Bacteroidetes species and shed light on functional aspects of a few hand-selected candidate sRNAs (reviewed in (9)). However, the phenotypic consequences of these sRNAs’ activities have largely remained obscure. Only for a single *Bacteroides* sRNA—termed MasB—could a loss-of-function phenotype be identified (10). Guided by TIS data, we found *masB*-deficient *B*. *thetaiotaomicron* mutants to have an enhanced tolerance of tetracyclines. In general terms, however, TIS—like any other random perturbation approach—biases against mutants of short genes, e.g. those of sRNAs. Thus, to systematically dissect the functional implications of these cellular regulators, tailored assays are required.

Acknowledging this need, we here introduced CRISPRi to sRNA research in *Bacteroides*. In light of the AT-rich nature of *Bacteroides* spp. genomes, we based our screen on the 5′-TTV-3′ PAM-recognizing nuclease Pb2Cas12a. The ensuing extended targeting space within the intergenic sRNA repertoire of *B*. *thetaiotaomicron* compared to that of SpCas9 allowed for more stringent spacer design rules, including “soft” gRNA parameters like optimal GC content and minimal risk of off-targeting. Our library design script is customizable, allowing users among others to select a desired PAM, and will hence be useful to construct libraries for other custom gene sets in virtually any bacterial species. As a compelling test case for the current study, we harnessed CRISPRi to assess the fitness of *B*. *thetaiotaomicron* sRNA knockdown mutants during bile stress. As a result, our screen proposed the sRNAs BatR and BTnc353 to reduce or enhance *B*. *thetaiotaomicron* bile stress resilience, respectively. While the BTnc353-associated bile phenotype needs to be further validated in the future, we confirmed BatR to indeed delay *B*. *thetaiotaomicron* outgrowth in the presence of bile salts. According to our target identification, BatR interacts *in vitro* with the 5′ regions of four mRNAs that encode for proteins involved in cell surface modulation. For two of these—*BT_0521* and *BT_1177*—BatR binding results in reduced transcript levels *in vivo*, while the steady-state levels of the two remaining candidates were not affected by the sRNA in this experimental condition (mid-exponential growth in rich medium). Obviously, this does not exclude the possibility that these candidates (*BT_1195*, *BT_3849*) are indeed *bona fide* BatR targets, whose regulation primarily manifests at the level of translation. In any case, it is worth noting that for efficient target binding, BatR seems to rely on 5′ end maturation. Such a processing-dependent liberation of an sRNA’s seed region is reminiscent of proteobacterial sRNAs, such as *Salmonella* ArcZ (35) and *Campylobacter* CJnc190 (36). While maturation of these sRNAs is catalyzed by RNase E or RNase III, respectively, the *Bacteroides* nuclease(s) responsible for BatR processing remain(s) elusive. In light of its associated loss-of-function phenotype, the relevance of its target genes for *Bacteroides* cell surface structure, and the implied, unusual sRNA biogenesis/maturation, BatR arises as a prime candidate for follow-up mechanistic and functional studies.

In the future, our CRISPRi approach may be used to screen for sRNA functions under a variety of further growth conditions. Of specific interest to *B*. *thetaiotaomicron* biology are experimental conditions mimicking the natural environment of these gut microbes, such as growth in the presence of carbon sources derived from the host or its diet (37). Beyond simplistic *in vitro* settings, the CRISPRi mutant library could be deployed for colonization experiments of cell culture or mouse models. Such *in vivo* applications typically involve growth over extended time spans and will therefore require careful planning of the experimental setup. For example, it will be essential to ensure continuously high concentrations of the nuclease inducer in order to maintain selective pressure throughout the experiment (note that constitutive expression of Cas nucleases is generally not an option due to the resulting toxic effects observed in multiple bacterial species (38–42)). This might be achieved through regularly replenishing fresh IPTG in the medium (for cell culture-based experiments) or drinking water (for animal models). Of note, in case of the latter, colonization with a small, predefined CRISPRi library likely overcomes the bottleneck effects often hampering *in vivo* screens based on genome-wide transposon insertion libraries composed of hundreds of thousands of individual mutants (14).

Beyond targeting noncoding RNAs, the resources described in this work add to the functional toolset available for *Bacteroides* spp. and shall be useful for various other applications. Small proteins, for example, are prevalent across the bacterial phylogenetic tree and have also been annotated in *Bacteroides* spp. (15, 43). Suffering from similar limitations associated with sRNAs, small protein mutants tend to be underrepresented in classical transposon libraries. In contrast, a targeted CRISPRi approach analogous to the one described here would allow researchers to systematically screen for small protein-associated phenotypes. Besides, our work lays the ground for future genetic interaction screens (44) in which CRISPRi constructs targeting defined gene sets would be introduced in a preexisting mutant background. Alternatively, our approach could be adapted for CRISPR activation (16), wherein a Cas nuclease is fused to a transcriptional activator, resulting in the specific upregulation of targeted genes. The functional insights gained from these and related studies would help improve our fundamental understanding of the bacterial members of the gut microbiota.

## MATERIAL AND METHODS

### Bacterial strains and culture conditions

Supplementary Table 1 lists the strains, plasmids, oligonucleotides, and media used in this work. *Bacteroides* strains were grown at 37°C in an anaerobic chamber (Coy Laboratory Products) using an anaerobic gas mix of 85% N_2_, 10% CO_2_, and 5% H_2_. Solid media plates were made of supplemented BHI. Liquid cultures were grown in either rich (TYG) or minimal medium supplemented with 0.5% glucose (MM-G) (Supplementary Table 1). For growth in rich media, a single colony of wild-type *B*. *thetaiotaomicron* VPI-5482 (AWS-001) was inoculated in 5 mL of TYG and incubated overnight (∼ 16 h) at 37°C. For growth in MM-G, a single colony was inoculated in 5 mL and incubated for 24 h. Subcultures were made from these starter cultures in a 1:100 dilution into fresh medium.

### PAM frequency determination

A custom Python script was used for determining the abundance of selected PAMs among the intergenic sRNAs or the entire VPI-5482 genome sequence. The script counts the number of occurrences of each PAM on either strand of the input sequences. The code is available at https://github.com/gprezza/CRISPRi_tools.

### Nuclease screening by in vitro transcription and translation (TXTL)

Cleavage assays were performed as described previously (31). The plasmids encoding Cas12a, T7 RNA polymerase, and the CRISPR array were added into 9 µL of MyTXTL master mix (Arbor Biosciences, Cat # MYtxtl-70-96-M) to the same final concentration of 4.17 nM and a total volume of 12 µL. The mix was then incubated at 29°C for 16 h. Then, 1 µL of the mix, 1 µL of the targeted deGFP-encoding plasmid (concentration 10 nM), and 1 µL of nuclease-free water were added into 9 µL of MyTXTL master mix. Aliquots of 5 µL were placed in the wells of a 96-well V-bottom plate (Corning Costar, Cat # 3357) and incubated at 29°C for 16 h in a Synergy Neo2 microplate reader (BioTek, USA) with kinetic reading every 3 min (excitation 485 nm, emission 528 nm, gain 60, light source – Xenon Flash).

### Western blotting

Bacterial cultures grown in either TYG or MM-G media with the indicated IPTG concentration were collected at the indicated OD/time points, pelleted, and resuspended in Laemmli buffer at a concentration of 0.1 OD/mL. 20 µL of each sample were separated by 7% Tris Acetate PAGE and blotted on PVDF membranes. Equal loading was verified by staining the membrane with Ponceau S solution. FLAG-tagged protein signals were detected with anti-FLAG (Sigma-Aldrich, F1804) in combination with anti-mouse secondary antibody (Invitrogen, 31430).

### Cloning of nucleases, gRNAs, and arrays

All clonings were carried on in a Gibson assembly-like reaction with the NEBuilder® HiFi DNA Assembly kit (NEB, E2621) and into plasmids derived from pMM704 or pMM731 (17), a gift from Timothy Lu & Christopher Voigt (Addgene plasmids #68891, http://n2t.net/addgene:68891, RRID:Addgene_68891 and #68896, http://n2t.net/addgene:68896, RRID:Addgene_68896). These vectors are based on the pNBU2 backbone, which integrates into the *Bacteroides* genome at either of the two *tRNA^Ser^* sites, ensuring preservation of a constant gene copy number of 1 across different transconjugants (PMID 10852890). The oligonucleotides used in the cloning reactions are indicated in Supplementary Table 1.

Plasmids AWP-001 and AWP-002 were derived from pMM704 (without luciferase) and pMM731 (with luciferase), respectively, by replacing the Cas9 sequencce with an *Asc*I-*Not*I insertion region and the gRNA with a cassette formed by an FnCas12a repeat, a *Sma*I site, and the T1 terminator sequence of the *E*. *coli rnpB* gene. *E*. *coli* codon-optimized, FLAG-tagged Pb2Cas12a and dFnCas12a were PCR-amplified from plasmids pCB841 (internal number: 132282) (22) and pCB873 (31), respectively. dPb2Cas12a was generated by site-directed mutagenesis. The dPb2Cas12a or dFnCas12a cassette was inserted between the *Asc*I-*Not*I sites of plasmid AWP-002, resulting in the plasmids AWP008 and AWP-006. AWP-029, the plasmid expressing dPb2Cas12a without the luciferase cassette, was cloned by inserting dPb2Cas12a into AWP-001.

For cloning gRNAs, oligonucleotides comprising the 20 nt spacer sequence flanked 5′ by (5′ATCTTTGCAGTAATTTCTACTGTTGTAGAT) and 3′ by (5′-CCGGCTTATCGGTCAGTTTCACCTGATTTA) ho-mology arms were joined to *Sma*I-linearized AWP-008 or AWP-029 in a 10 μL reaction in a 20:1 oligonucleotide:vector ratio. Single-spacer arrays were similarly constructed by inserting 30 nt spacers into either AWP-009 (*Francisella* repeats; 5′ arm: 5′TTAAATAATTTCTACTGTTGTAGAT, 3′ arm: 5′-GTCTAAGAACTTTAAATAATTTGTC) or AWP-010 (*Prevotella* repeats; 5′ arm: 5′-TTATTTAATTTCTACTATTGTAGAT, 3′ arm: 5′GGCTATAAAGCTTATTTAATTTCTA). Three-spacer arrays were cloned through CRATES (PMID: 31270316).

All cloned plasmids were introduced in the donor *E*. *coli* S17-1 λpir strain. Transformants were conjugated with wild-type *B*. *thetaiotaomicron* and transconjugants counterselected on plates containing gentamicin and erythromycin.

### Luciferase assay

Endpoint measurements of Nanoluc activity were performed as previously described (17) with minor modifications. An overnight culture of a Nanolucand dPb2Cas12a-expressing strain (containing a plasmid derived from AWP-008, AWP-009 or AWP-010) was subcultured 1:100 in fresh medium containing 250 μM IPTG to induce expression of the nuclease. Cultures were removed from the anaerobic chamber and put on ice at OD_600_ 0.3-0.4 (∼5 h). An aliquot of the culture was stored on ice for OD measurement at the end of the protocol. 500 µL of each sample were pelleted, washed once with PBS, and pelleted again, after which cells were lysed by resuspending in 50 μL of BugBuster reagent (Merck, 71456) and incubation under constant rotation for 10 min at room temperature. The lysate was cleared by centrifugation at 15,000 g for 15 min at 4°C and 45 μL of the supernatant were mixed with an equal volume of the NanoLuc Reagent (Promega N1110) prepared according to the manufacturer’s instructions. After incubation for 5 min at 26°C, light emission of the sample was measured in 2 x 40 μL technical replicates on a Tecan infinite 200 PRO plate reader. Luciferase measurements were normalized to OD_600_ values of 100 μL of the starting cultures, as measured in parallel on the infinite 200 PRO plate reader.

### RNA extraction, northern blotting, qRT-PCR

Bacterial cultures were grown until the indicated OD and RNA was extracted as previously described (20). Briefly, 4 OD equivalents were collected in a tube, mixed with 20% volume of stop mix (45) (95% ethanol, 5% water saturated phenol, pH >7.0) and snap-frozen in liquid nitrogen. After pelleting, cells were lysed with lysozyme (0.5 mg/mL) and 10% SDS in TE buffer. After addition of NaOAc (0.3 M final concentration), phenol (pH 4.5-5) was mixed at 1:1 volume and the samples were incubated for 6 min at 65°C, followed by addition of 1:1 volume of chloroform. After separation by centrifugation, nucleic acids in the aqueous phase were precipitated by adding 2 volumes of ethanol:3 M NaOAc (30:1 ratio) and incubating overnight at -80°C. After centrifugation, the pellet was resuspended in water and 40 μg of total RNA was cleared of contaminating DNA through treatment with 5 U of DNase I (Thermo Fisher Scientific #EN0521). RNA was again purified with a phenol-chloroform extraction followed by ethanol-NaOAc precipitation.

For northern blotting, 5 μg of total RNA was mixed with 1 volume of loading dye (95% formamide, 18 mM EDTA, 0.025% SDS, 0.025% xylene cyanol, 0.025% bromophenol blue) and denatured for 5 min at 95°C followed by 5 min incubation on ice. RNA was separated on a denaturing 6% polyacrylamide-7 M urea gel and electroblotted onto Hybond XL membranes (Amersham). Blotted membranes were probed with ^32^P-labeled gene-specific oligonucleotides (Supplementary Table 1).

qRT-PCR was performed in a one-step reaction that includes reverse transcription with Takyon UF-NSMT-B0701 master mix and UF-RTAD-D0701 One-Step Kit Converter. The final volume of each reaction was 10 μL, containing 20 ng total RNA, 5 μL of qPCR mastermix, 0.1 μL of each primer (10 μM), and 0.1 μL of reverse transcriptase. Three technical replicates per sample were measured.

### Design and cloning of the CRISPRi library

The script used to design the library is available at https://github.com/gprezza/CRISPRi_tools. Version v0.1.0 was used, with options *-a –nt*. A description of its criteria for selecting spacers and arrays can be found in the “Automated design of gRNAs against the full suite of intergenic *B*. *thetaiotaomicron* sRNAs” Results section. A Cas12a array is considered as properly folded if the probability of forming all three repeat hairpins is >20%, as predicted by RNAfold (46). The library was designed against a prior annotation of intergenic sRNAs, and some targeted sRNAs have since been reassigned to other categories (10). These sRNAs were filtered out during data analysis. The designed guide sequences are available in Supplementary Table 2. The script provides oligonucleotides for cloning Cas12a three-spacer arrays with CRATES (31) reactions into AWP-031, which was derived from AWP-029 by replacing the gRNA cassette with a GFP dropout system flanked by *Bsm*BI sites at both ends as in the original CRATES dropout construct. Restriction digestion leaves the overhangs CCTC and AACG at the 5′ and 3′ end, respectively. These overhang sequences are needed for CRATES and were taken from the “set 3” group of 25 unique four-base overhangs that were predicted to have > 95% fidelity in Gibson-like reactions in a previous study (47). The remaining 23 overhangs from the set allow cloning up to 23 arrays in a single CRATES reaction, in which each three-spacer array is generated from a set of six partially overlapping oligonucleotides (31). The final assembled array is flanked by the CCTC and AACG overhangs and contains internally one of the 23 overhangs, arranged so that only the full array comprising all three spacers can be formed. The script divided the set of 91 designed arrays in three sets of 23 arrays each (138 oligonucleotides) and a fourth set of 22 arrays (132 oligonucleotides).

These four pools were ordered as oPools^TM^ oligo pools (IDT) and cloned into AWP-031 in four distinct CRATES reactions, with each containing all 138 or 132 oligonucleotides of the pool. The reactions were carried out similarly to the original protocol (31), with some modifications. Each oligonucleotide pool was phosphorylated at 37°C for 1 h, followed by a 65°C incubation for 20 min in a 40 μL reaction containing 5 μL of the pool (350 ng/µL, ∼10 μM), 1.5 μL T4 Polynucleotide Kinase (Thermo #EK0031), 4 μL PNK buffer A, 4 μL of 10 mM ATP, and 25.5 μL of water. Phosphorylated oligonucleotide pools were annealed by incubating for 5 min at 95°C, followed by cooling down to 85°C in steps of 30 s/degree, then to 65°C at 1 min/degree, and finally to 15°C at 30 s/degree. After this step, the nicks in the annealed fragments were repaired in a 50 μL reaction kept at 16°C for 5 h, followed by 65°C for 10 min and containing 40 μL of the annealed oligonucleotide pool, 2 μL of 10 mM ATP, 2.5 μL T4 DNA ligase (NEB #M0202), 0.55 μL PNK buffer A (Thermo), and 4.95 μL of water. We noticed that the assembled fragments comprised either the full-length three-spacer array or some shorter, misassembled sequences. We therefore gel-extracted the band corresponding to the size of the correctly assembled arrays (202 bp) and ligated these into already *Bsm*BI-digested AWP-031 in a reaction containing 80 ng digested AWP-031, 30 ng gel-extracted arrays, 1 μL T4 ligase buffer, 0.5 μL T4 DNA ligase, 0.5 μL *Bsm*BI, and water to 10 μL. The ligation was incubated in a thermocycler using the following program: 45 cycles of alternating digestion and ligation (42°C for 2 min, 16°C for 5 min) followed by a final digestion (55°C for 20 min) and an heat-inactivation step (80°C for 10 min).

Oligonucleotides for cloning gRNAs into AWP-029 are also provided by the script, comprising the 20 nt spacer sequence flanked by a 5′ (5′-ATCTTTGCAGTAATTTCTACTGTTGTAGAT) and a 3′ (5′CCGGCTTATCGGTCAGTTTCACCTGATTTA) homology arm. The oligonucleotides containing the 557 spacers designed by the script were ordered as two oPools^TM^ oligo pools (IDT) comprising 278 and 279 spacers each. The pools were cloned into *Sma*I-linearized AWP-029 with the NEBuilder® HiFi DNA Assembly kit (NEB, E2621) in two separate reactions as described above for the generation of single-gRNA mutants.

Ligated gRNA and array pools were separately transformed into the *E*. *coli* S17-1 λpir strain. We collected and pooled 1-2x10^4^ colonies per transformation reaction and generated glycerol stocks. We inoculated each transformant pool in LB medium supplemented with ampicillin at a starting concentration of 0.04 OD, let grow until early exponential phase (∼ OD 0.5), and mixed equal volumes of each array pool and each gRNA pool. We then mixed the array and gRNA *E*. *coli* cultures with wild-type *B*. *thetaiotaomicron* in early exponential phase (∼ OD 0.7) at a 1:4:10 ratio, plated 10 OD of the mix on BHIS plates, and incubated aerobically overnight at 37°C. The following day, we resuspended the cell lawn in 1 mL of PBS and plated 100 μL aliquots of 10^-1^ and 10^-2^ dilutions on BHIS plates containing gentamicin (100 µg/mL) and erythromycin (12.5 μg/mL). We collected ∼3x10^5^ colonies and inoculated 70 mL of TYG supplemented with the same antibiotics at a starting OD of 0.08 OD. After the culture reached stationary phase (OD 4.2; ∼24 h of incubation at 37°C), it was harvested and a glycerol stock was prepared. Three aliquots of the glycerol stock were processed for gDNA extraction and sequencing as described below. We considered mutants with >10 read counts in all biological replicates as present in the final library.

### CRISPRi screen under bile salt stress

We spread an aliquot of the glycerol stock of the guide library on a BHIS plate and incubated for 2-3 d at 37°C. The day before the experiment, making sure to avoid single colonies, we harvested part of the cell lawn and used it to start an overnight culture in TYG medium at a starting OD of 0.05 OD/mL. We then diluted this input culture 1:100 into fresh TYG medium containing 250 μM of IPTG, with or without 0.5 mg/mL of bile salts (Fluka #48305). We incubated the subculture at 37°C until it reached the indicated OD, at which point we took a 2 mL aliquot, pelleted it, discarded the supernatant, and stored the bacterial samples at -20°C until further processing.

Bacterial pellets were resuspended in 100 μL of PBS containing 250 μg lysozyme, mixed with 100 μL lysis buffer (100 mM Tris-HCl pH 8.5, 200 mM NaCl, 0.2% SDS, 5 mM EDTA), and lysed by incubating at 37°C for 5 min. Proteins were then degraded by addition of 1 μL of Proteinase K (NEB #P8107) and incubation at 56°C for 30 min, following which RNA was digested with 0.5 μL of RNase A (Thermo #EN0531) and incubation at 56°C for 5 min. The remaining gDNA was purified from each sample via phenol-chloroform extraction, followed by ethanol precipitation.

The genomic regions coding for gRNAs and arrays were PCR-amplified in a 50 µL reaction containing 100 ng gDNA with oligonucleotides AWO-577/AWO-578 (annealing temperature: 55°C) and AWO-575/AWO-576 (annealing temperature: 63°C).

### Sequencing and data analysis for the CRISPRi mutant fitness screen

Array and gRNA amplicons from the same gDNA sample were mixed at a 3:1 molar ratio and sequenced on an Illumina NextSeq 500 or NextSeq 2000 platform in paired-end mode, 150 bp reads. Data are available at the National Center for Biotechnology Information Gene Expression Omnibus database (https://www.ncbi.nlm.nih.gov/geo) under the accession number GSE235620. Sequenced reads were adapterand qualitytrimmed with BBDuk, using the following parameters: *k=13 mink=5 rcomp=t literal=ACACTCTTTCCCTACACGACGCTCTTCCGATCT,GTGACTGGAGTTCAGACGTGTGCTCTTCCGATCT hdist=1ktrim=r minlen=100 tbo*. Trimmed reads were merged with BBMerge, using parameters *mininsert=120 maxloose=t*. The abundance of each CRISPRi mutant within the merged reads was determined with a custom Python script (https://github.com/gprezza/CRISPRi_tools). Sequences of gRNAs and assembled arrays used for read mapping are available in Supplementary Table 3.

Differential mutant abundance analysis was performed with the R package edgeR (3.32.1), separately for gRNAs and arrays. In each case, mutants with low abundance were filtered out with the filterByExpr function and undesired batch effects were removed with the RUVs function of the RUVSeq package (48). Normalization of the libraries was done with the *betweenLaneNormalization* function (“upper” method) of the EDASeq package. Fitness associated with the inhibition of each sRNA was calculated by averaging the fold-change of all mutants targeting the same sRNA, while their p-values were combined with Fisher’s method. Differential mutant abundance and sRNA fitness scores can be found in Supplementary Table 4.

### sRNA mutant strain generation

The *batR* gene was deleted from the *B*. *thetaiotaomicron* VPI-5482 genome as previously described (49). Briefly, we cloned ∼750 bp of the regions upstream and downstream of the sRNA into the multiple cloning site of pSIE1 (a gift from Andrew Goodman, Addgene plasmid #136355; http://n2t.net/addgene:136355; RRID:Addgene_136355). The plasmid was then conjugated into *B*. *thetaiotaomicron*. Transconjugant colonies were grown overnight in TYG medium and streaked on BHIS plates containing anhydrotetracycline. Mutants that had correctly excised the plasmid backbone were identified by colony PCR and the deletion and absence of unwanted mutations surrounding it were confirmed by Sanger sequencing.

### B. thetaiotaomicron growth curves

Overnight cultures of a single colony of the indicated strains were diluted 1:100 into fresh TYG medium with or without 0.05 mg/mL of bile salts (Fluka #48305) and aliquoted in four 200 µL technical replicates in a transparent 96-well plate. Growth at 37°C was recorded by measuring absorbance (600 nm) in 20 min intervals with 10 s shaking before each measurement.

### sRNA homolog search, target prediction and synteny analysis

The sequence of BatR, including 50 nucleotides upstream of the sRNA’s 5′ end to comprise the promoter, was searched with blastn (7 nt word size) (50) against the refseq_representative_genomes database filtered for Bacteroidetes (taxid 976). Sequences with <50% query cover were discarded and alignments were made with MAFFT (51). sRNA target prediction was done using the CopraRNA webserver (34) with default options, except for “nt up”, which was set to 100 nt.

Synteny analysis was done on the region 5000 nt up- and down-stream of each BatR homolog (154 nt isoform). The annotated subgenomic regions were retrieved from the NCBI Nucleotide database and the synteny figure was generated with clinker (52).

### In vitro transcription and 5′ end labelling

RNAs were transcribed with the MEGAscript T7 kit (Ambion) according to the manufacturer’s manual from templates prepared by PCR amplification of the respective genomic regions, with a forward primer containing the T7 promoter sequence. After transcription, samples were incubated with 1U TURBO DNase, Thermo Scientific for 15 min at 37°C. The RNA products were then separated on a 7 M urea 6% PAGE and bands of the correct size were cut out. RNA was eluted from the bands by overnight incubation at 4°C and 1,000 rpm shaking in 750 μL of RNA elution buffer (0.1 M NaAc, 0.1% SDS, 10 mM EDTA), following which the supernatant was subjected to phenol-chloroform extraction and ethanol precipitation to obtain the purified, *in vitro* transcribed RNAs. 50 pmol of each RNA were dephosphorylated by incubation for 1 h at 37°C with 25 U of calf intestine alkaline phosphatase (NEB) in a final volume of 50 µL, following purification by phenol-chloroform extraction and ethanol precipitation. 20 pmol of the obtained RNA were then radiolabeled by incubation with 1 U of Polynucleotide Kinase (NEB) and 20 μCi of ^32^P-γATP for 1 h at 37°C in a 20 μL reaction volume. Labeled RNA was purified on a G-50 column (GE Healthcare), following separation in and extraction from a polyacrylamide gel as described above.

### Electrophoretic mobility shift assay (EMSA)

EMSAs were performed by incubating 4 nM of 5′ radiolabeled BatR with increasing concentrations (0, 3.9, 7.8, 15.6, 31.3, 62.5, 125, 250, 500 nM) of *in vitro*-transcribed mRNA fragments. Before incubation, the mRNAs were denatured by heating at 95°C for 1 min, followed by incubation on ice. The two RNA partners were then mixed in 1× structure buffer (10 mM Tris–HCl pH 7.0, 0.1 M KCl, 10 mM MgCl_2_) containing 0.1 μg/μL of yeast RNA (Ambion) and incubated at 37°C for 1 h. Immediately prior loading, reactions were stopped by adding 3 μL of 5× loading dye (0.5× TBE, 50% glycerol, 0.2% xylene cyanol, 0.2% bromophenol blue). Samples were separated on a native 6% polyacrylamide gel, run at 4°C in 0.5% TBE buffer. After 3 h the run was stopped, gels were dried, and exposed on phosphor screens.

The dissociation constant (Kd) was calculated by fitting the values of the fraction of bound sRNAs (F_bound_) to an exponential curve of formula F_bound_ = B_max_ * mRNA_conc_ / (Kd + mRNA_conc_), where B_max_ is the maximum specific binding and mRNA_conc_ the concentration of mRNA fragment.

## ACKNOWLEDGEMENTS

This work was supported by a European Research Council (ERC) Starting Grant (101040214 to A.J.W.) and an ERC Consolidator Award (865973 to C.L.B.).

## SUPPLEMENTARY FILES

**Supplementary Figure S1.**
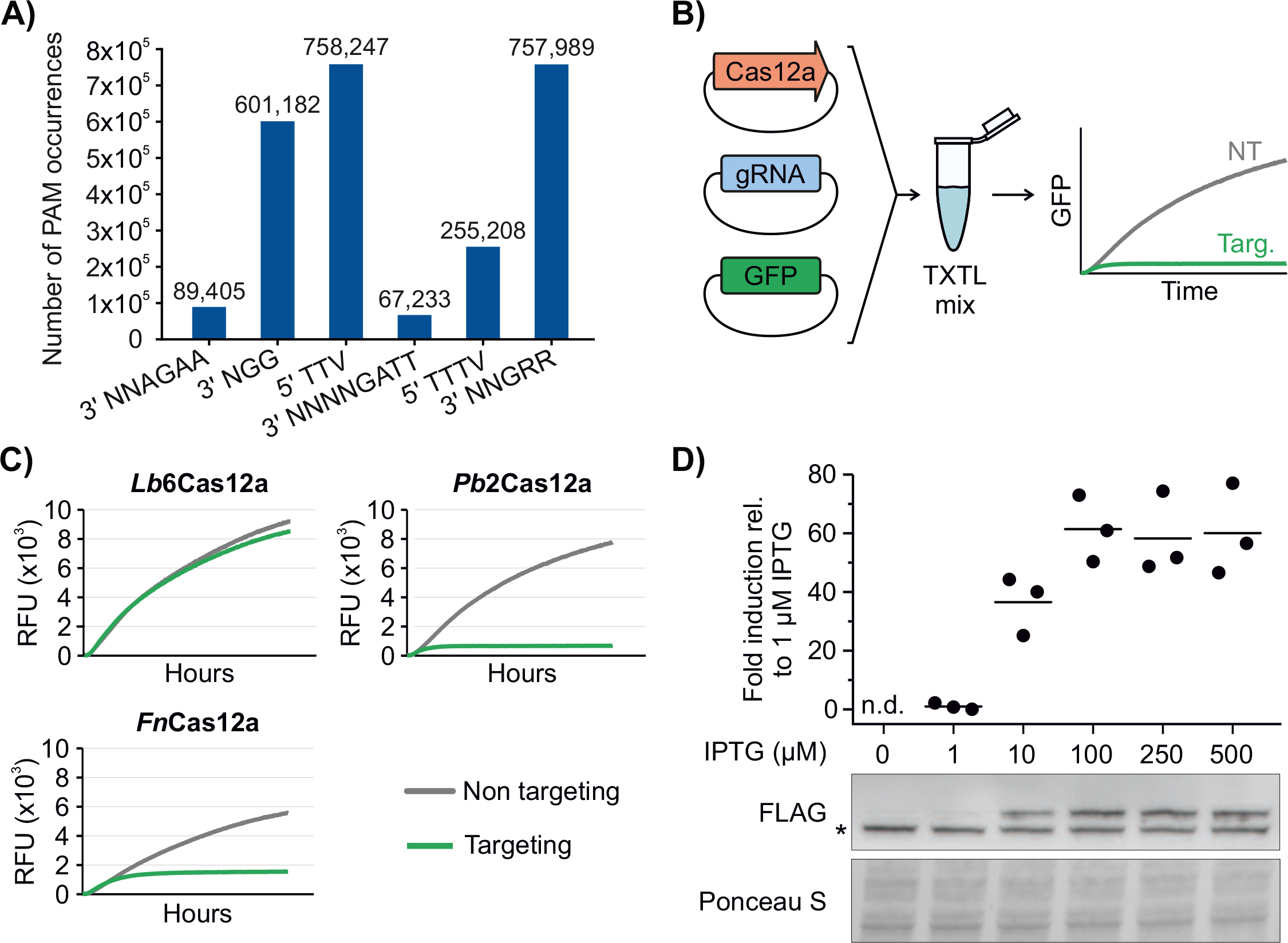
A) Frequency of different PAMs within the entire genome of *B*. *thetaiotaomicron* VPI-5482. Related to Fig. 1A. B) Scheme of the TXTL assay. A plasmid encoding GFP was co-expressed in a TXTL reaction with a Cas12a ortholog and a CRISPR array comprising either a gRNA targeting GFP or a nontargeting control. GFP fluorescence emission was recorded over time. A lower emission of the targeting array (Targ, green) versus the non-targeting one (NT, grey) indicate effective cleavage of the GFP coding sequence by Cas12a. C) Plasmid clearance assay using three Cas12a orthologs. Relative fluorescence units (RFU) are plotted over time for each separate reaction (n=1). D) Identification of the optimal IPTG concentration for induced expression of dPb2Cas12a. An overnight culture of a strain encoding dPb2Cas12a expressed under an IPTG-inducible promoter was subcultured 1:100 in fresh medium containing the indicated IPTG concentration. Top: quantification of dPb2Cas12a levels in three biological replicates, with a line representing the mean value. Bottom: representative western blot image and Ponceau S staining as loading control. An asterisk next to the blot indicates an unspecific band.

**Supplementary Figure S2.**
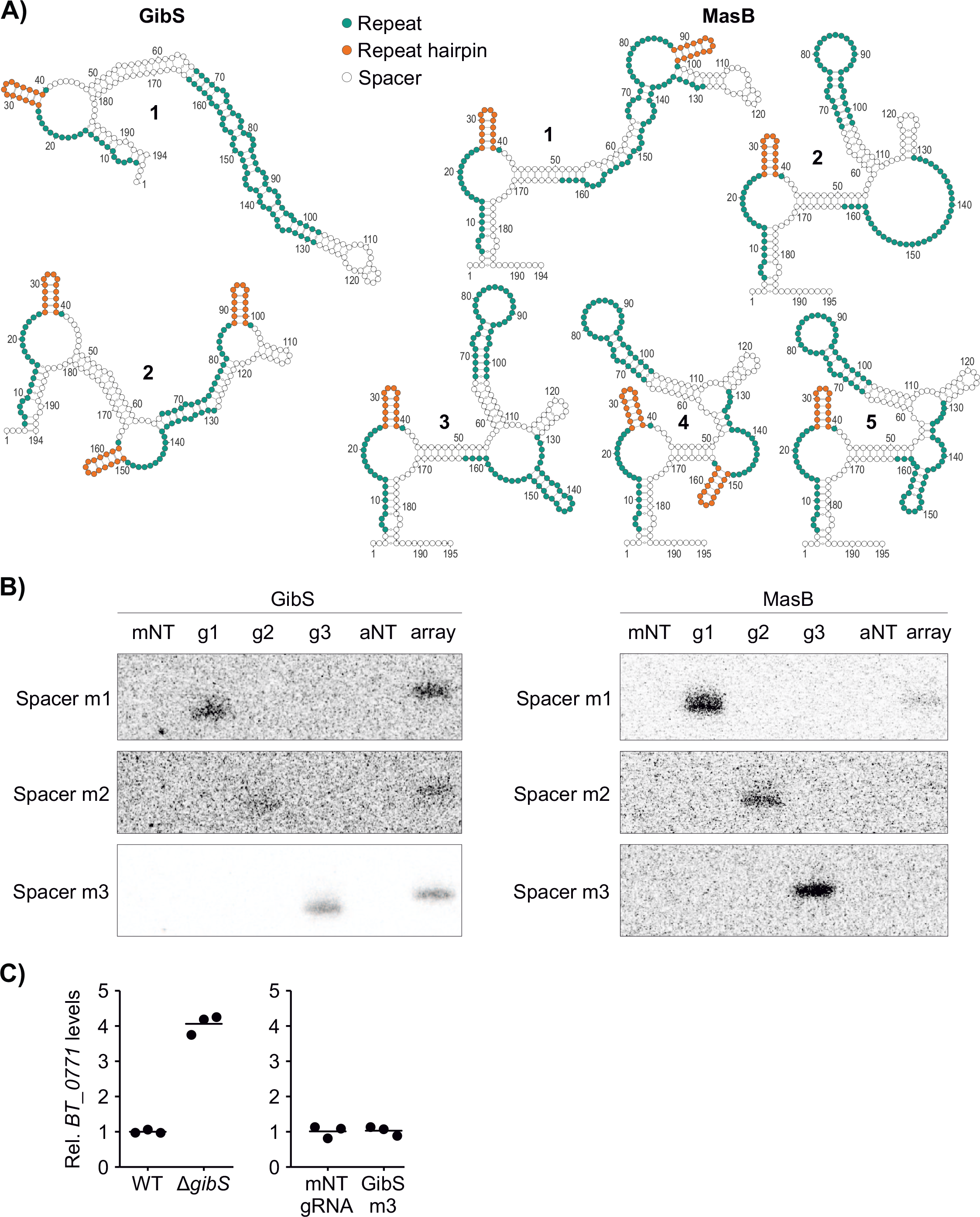
A) Predictions of the secondary structure of the GibS- and MasB-targeting arrays. The five structures with the lowest minimal free energy as predicted by mfold are shown, with a number next to each structure marking the rank position from lowest to highest predicted minimal free energy. Only two structures were predicted for the GibS array. Repeat sequences are highlighted in green, correctly formed terminal hairpins in orange. B) Detection by northern blotting of processed gRNA levels in strains expressing stand-alone gRNAs or the same spacers encoded in an array. For the corresponding 5S rRNA signal (loading control), see Fig. 3A. The probes used in the experiment anneal to each spacer sequence. Images are representative of two biological replicates. C) qRT-PCR measurements of *BT_0771* levels in Δ*tdk* (WT), Δ*tdk*Δ*gibS* (Δ*gibS*) or in bacteria harboring the CRISPRi vector with a non-targeting (gNT gRNA) or GibS-targeting (GibS g3) gRNA. Values shown are relative to Δ*tdk* or the non-targeting gRNA, respectively. A line represents the mean value.

**Supplementary Figure S3.**
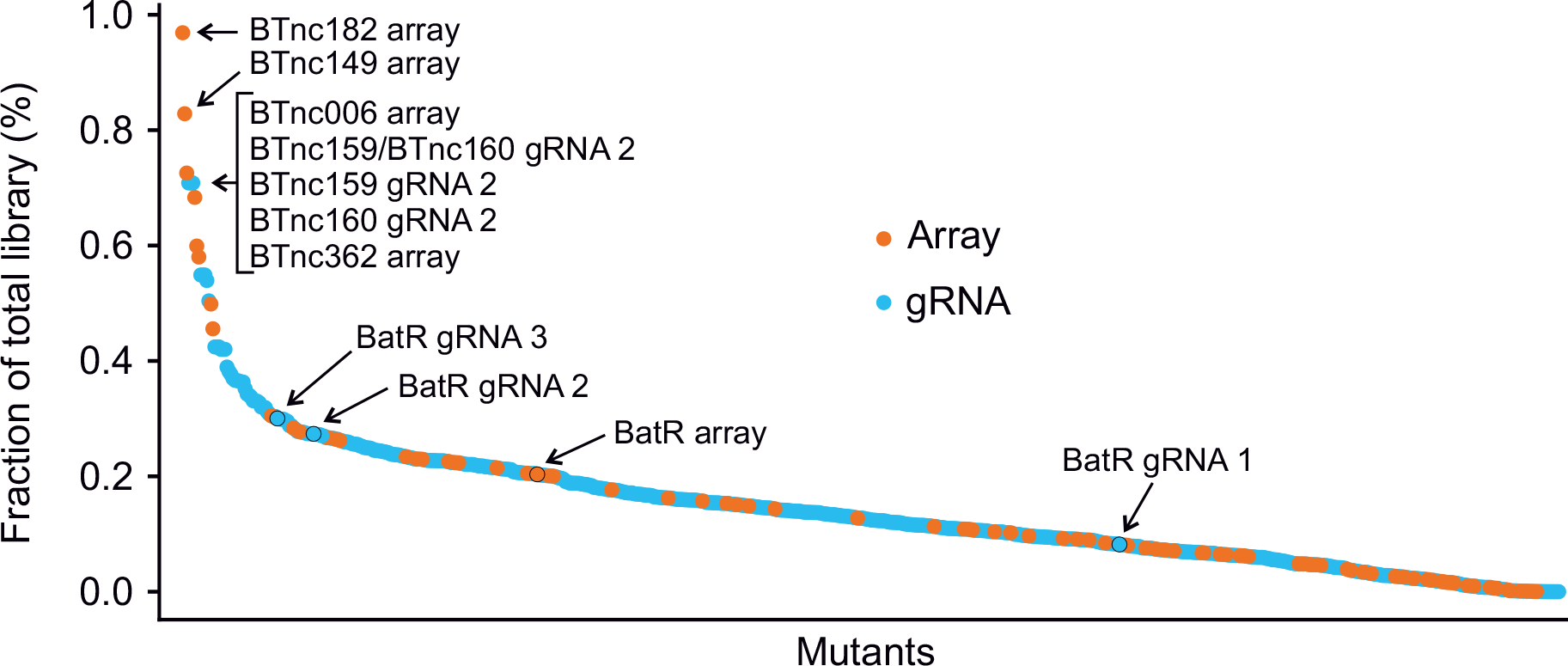
Distribution of each CRISPRi mutant in the guide library as fraction of the total mutants. The most abundant mutants, as well as those targeting BatR are labeled.

**Supplementary Figure S4.**
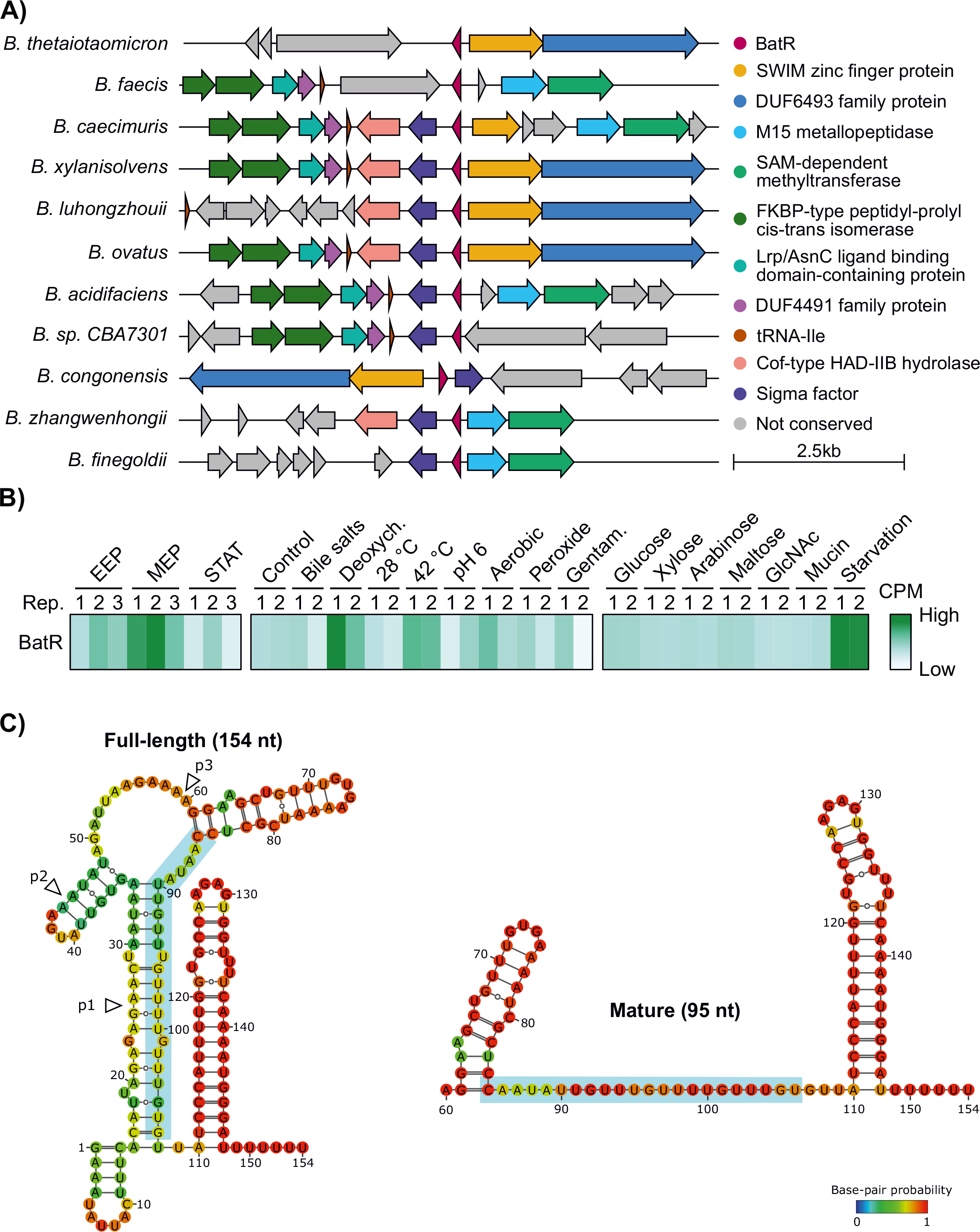
A) Synteny analysis of BatR homologs across *Bacteroides* spp. For each strain, the 5,000 nts upstream and downstream of the *batR* gene are plotted. Genes within these regions are represented as arrows and colored according to their function. B) Abundance (z-score of CPM values) of BatR during growth in rich medium (EEP: early exponential phase, MEP: mid-exponential phase, STAT: stationary phase), after exposure to different stressors or while feeding on defined carbon sources. C) Predicted secondary structure of full-length BatR or its shortest detected isoform (see Fig. 6B). Putative processing sites are indicated by white arrowheads and the seed region by a light blue background box.

**Supplementary Figure S5.**
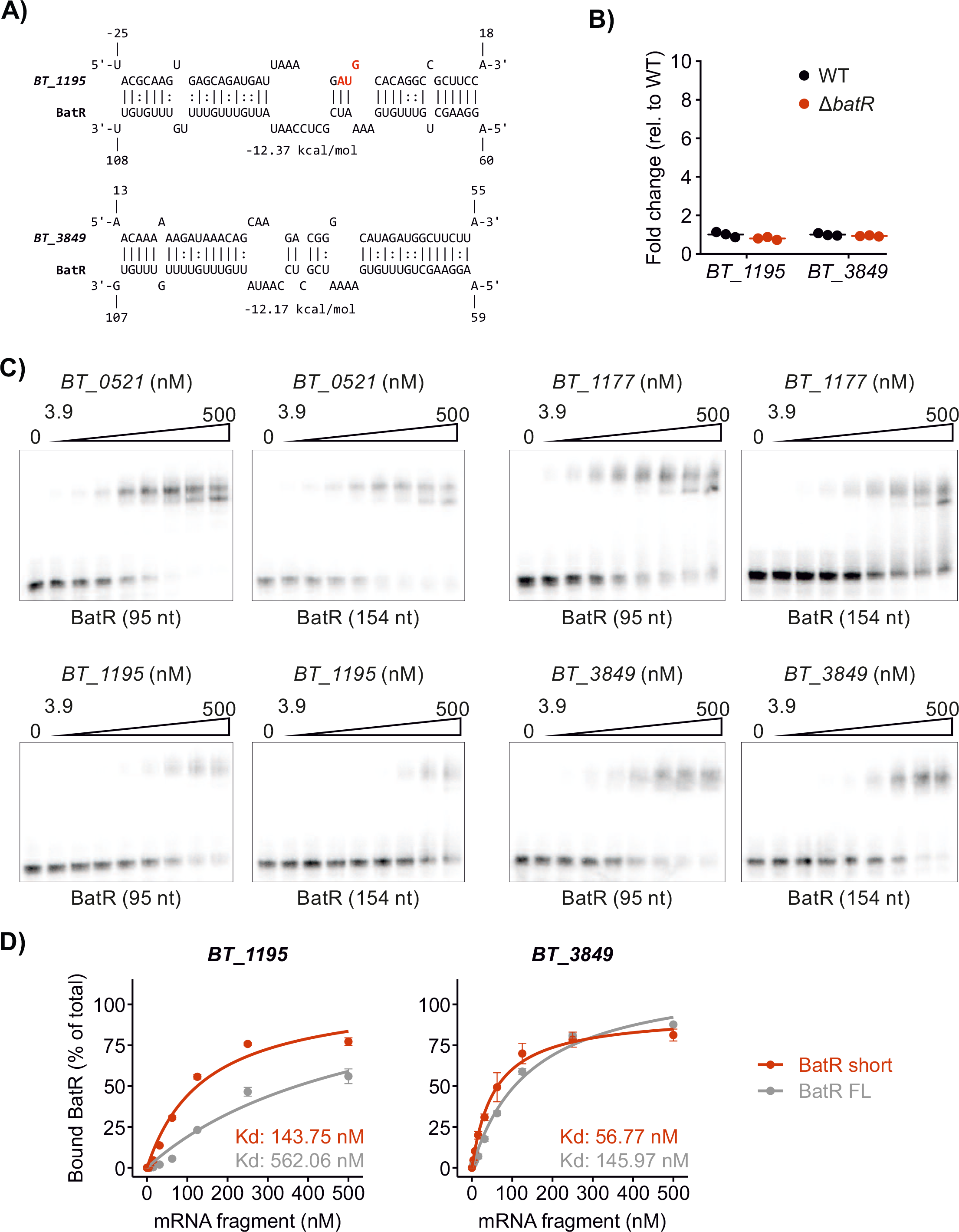
A) Predicted interaction between BatR and two putative mRNA targets.

B) qRT-PCR measurement of the levels of *BT_1195* and *BT_3849* in the wild-type and Δ*batR* strains.

C) EMSA of BatR and 5′ fragments of four predicted mRNA targets encompassing the predicted interaction sites (see Fig. 6C and Supplementary Fig. S5A). BatR at a constant (4 nM) concentration was mixed with increasing (0 to 500 nM) concentrations of each mRNA fragment. Each gel is representative of two replicates.

D) Quantification of the fraction of bound BatR at increasing concentrations of *BT_1195* (left) or *BT_3849* (right). Error bars represent standard deviation of two biological replicates.

Supplementary Table 1

Oligonucleotides, plasmids, strains, and media used in this study.

Supplementary Table 2

Oligonucleotides used for cloning the CRISPRi library.

Supplementary Table 3

Assembled sequences of all mutants in the CRISPRi library in FASTA format.

Supplementary Table 4

CRISPRi fold change data upon bile stress per each mutant (first sheet) or sRNA (second sheet).

Supplementary Table 5

CopraRNA target predictions for BatR.

## REFERENCES

1. Human Microbiome Project C (2012) Structure, function and diversity of the healthy human microbiome. Nature 486(7402):207–214.

2. Zafar H & Saier MH, Jr. (2021) Gut Bacteroides species in health and disease. Gut Microbes 13(1):1–20.

3. Wexler HM (2007) Bacteroides: the good, the bad, and the nitty-gritty. Clin Microbiol Rev 20(4):593–621.

4. Wexler AG & Goodman AL (2017) An insider’s perspective: Bacteroides as a window into the microbiome. Nat Microbiol 2:17026.

5. Gottesman S & Storz G (2011) Bacterial small RNA regulators: versatile roles and rapidly evolving variations. Cold Spring Harb Perspect Biol 3(12).

6. Wagner EGH & Romby P (2015) Small RNAs in bacteria and archaea: who they are, what they do, and how they do it. Advances in genetics 90:133–208.

7. Papenfort K & Melamed S (2023) Small RNAs, Large Networks: Posttranscriptional Regulons in Gram-Negative Bacteria. Annu Rev Microbiol.

8. Holmqvist E & Wagner EGH (2017) Impact of bacterial sRNAs in stress responses. Biochem Soc Trans 45(6):1203–1212.

9. Ryan D, Prezza G, & Westermann AJ (2020) An RNA-centric view on gut Bacteroidetes. Biol Chem 402(1):55–72.

10. Ryan D, et al. (2023) An integrated transcriptomics-functional genomics approach reveals a small RNA that modulates Bacteroides thetaiotaomicron sensitivity to tetracyclines. *bioRxiv*.

11. Goodman AL, et al. (2009) Identifying genetic determinants needed to establish a human gut symbiont in its habitat. Cell Host Microbe 6(3):279–289.

12. Liu H, et al. (2021) Functional genetics of human gut commensal Bacteroides thetaiotaomicron reveals metabolic requirements for growth across environments. Cell Rep 34(9):108789.

13. Chao MC, Abel S, Davis BM, & Waldor MK (2016) The design and analysis of transposon insertion sequencing experiments. Nature reviews. Microbiology 14(2):119–128.

14. Cain AK, et al. (2020) A decade of advances in transposon-insertion sequencing. Nat Rev Genet 21(9):526–540.

15. Gray T, Storz G, & Papenfort K (2022) Small Proteins; Big Questions. J Bacteriol 204(1):e0034121.

16. Todor H, Silvis MR, Osadnik H, & Gross CA (2021) Bacterial CRISPR screens for gene function. Curr Opin Microbiol 59:102–109.

17. Mimee M, Tucker AC, Voigt CA, & Lu TK (2015) Programming a Human Commensal Bacterium, Bacteroides thetaiotaomicron, to Sense and Respond to Stimuli in the Murine Gut Microbiota. Cell Syst 1(1):62–71.

18. Wang T, et al. (2018) Pooled CRISPR interference screening enables genome-scale functional genomics study in bacteria with superior performance. Nature communications 9(1):2475.

19. Rishi HS, et al. (2020) Systematic genome-wide querying of coding and non-coding functional elements in E. coli using CRISPRi.2020.2003.2004.975888.

20. Ryan D, Jenniches L, Reichardt S, Barquist L, & Westermann AJ (2020) A high-resolution transcriptome map identifies small RNA regulation of metabolism in the gut microbe Bacteroides thetaiotaomicron. Nature communications 11(1):3557.

21. Deveau H, et al. (2008) Phage response to CRISPR-encoded resistance in Streptococcus thermophilus. J Bacteriol 190(4):1390–1400.

22. Marshall R, et al. (2018) Rapid and Scalable Characterization of CRISPR Technologies Using an E. coli Cell-Free Transcription-Translation System. Mol Cell 69(1):146–157 e143.

23. Zhang Y, et al. (2013) Processing-independent CRISPR RNAs limit natural transformation in Neisseria meningitidis. Mol Cell 50(4):488–503.

24. Zetsche B, et al. (2015) Cpf1 is a single RNA-guided endonuclease of a class 2 CRISPR-Cas system. Cell 163(3):759–771.

25. Kleinstiver BP, et al. (2015) Broadening the targeting range of Staphylococcus aureus CRISPRCas9 by modifying PAM recognition. Nat Biotechnol 33(12):1293–1298.

26. Garamella J, Marshall R, Rustad M, & Noireaux V (2016) The All E. coli TX-TL Toolbox 2.0: A Platform for Cell-Free Synthetic Biology. ACS Synth Biol 5(4):344–355.

27. Zhang X, et al. (2017) Multiplex gene regulation by CRISPR-ddCpf1. Cell Discov 3:17018.

28. Miao C, Zhao H, Qian L, & Lou C (2019) Systematically investigating the key features of the DNase deactivated Cpf1 for tunable transcription regulation in prokaryotic cells. Synth Syst Biotechnol 4(1):1–9.

29. Zhang JL, et al. (2018) Gene repression via multiplex gRNA strategy in Y. lipolytica. Microb Cell Fact 17(1):62.

30. Fonfara I, Richter H, Bratovic M, Le Rhun A, & Charpentier E (2016) The CRISPR-associated DNA-cleaving enzyme Cpf1 also processes precursor CRISPR RNA. Nature 532(7600):517–521.

31. Liao C, et al. (2019) Modular one-pot assembly of CRISPR arrays enables library generation and reveals factors influencing crRNA biogenesis. Nature communications 10(1):2948.

32. Zuker M (2003) Mfold web server for nucleic acid folding and hybridization prediction. Nucleic Acids Res 31(13):3406–3415.

33. Collins SL, Stine JG, Bisanz JE, Okafor CD, & Patterson AD (2023) Bile acids and the gut microbiota: metabolic interactions and impacts on disease. Nature reviews. Microbiology 21(4):236–247.

34. Wright PR, et al. (2014) CopraRNA and IntaRNA: predicting small RNA targets, networks and interaction domains. Nucleic Acids Res 42(Web Server issue):W119-123.

35. Chao Y, et al. (2017) In Vivo Cleavage Map Illuminates the Central Role of RNase E in Coding and Non-coding RNA Pathways. Mol Cell 65(1):39–51.

36. Svensson SL & Sharma CM (2021) RNase III-mediated processing of a trans-acting bacterial sRNA and its cis-encoded antagonist. Elife 10.

37. Glowacki RWP & Martens EC (2020) If you eat it, or secrete it, they will grow: the expanding list of nutrients utilized by human gut bacteria. J Bacteriol 203(9).

38. Lee YJ, Hoynes-O’Connor A, Leong MC, & Moon TS (2016) Programmable control of bacterial gene expression with the combined CRISPR and antisense RNA system. Nucleic Acids Res 44(5):2462–2473.

39. Wendt KE, Ungerer J, Cobb RE, Zhao H, & Pakrasi HB (2016) CRISPR/Cas9 mediated targeted mutagenesis of the fast growing cyanobacterium Synechococcus elongatus UTEX 2973. Microb Cell Fact 15(1):115.

40. Rock JM, et al. (2017) Programmable transcriptional repression in mycobacteria using an orthogonal CRISPR interference platform. Nat Microbiol 2:16274.

41. Jiang Y, et al. (2017) CRISPR-Cpf1 assisted genome editing of Corynebacterium glutamicum. Nature communications 8:15179.

42. Cui L, et al. (2018) A CRISPRi screen in E. coli reveals sequence-specific toxicity of dCas9. Nature communications 9(1):1912.

43. Sberro H, et al. (2019) Large-Scale Analyses of Human Microbiomes Reveal Thousands of Small, Novel Genes. Cell 178(5):1245–1259 e1214.

44. Wong AS, Choi GC, & Lu TK (2016) Deciphering Combinatorial Genetics. Annu Rev Genet 50:515–538.

45. Tedin K & Blasi U (1996) The RNA chain elongation rate of the lambda late mRNA is unaffected by high levels of ppGpp in the absence of amino acid starvation. J Biol Chem 271(30):17675–17686.

46. Lorenz R, et al. (2011) ViennaRNA Package 2.0. Algorithms Mol Biol 6:26.

47. Potapov V, et al. (2018) Comprehensive Profiling of Four Base Overhang Ligation Fidelity by T4 DNA Ligase and Application to DNA Assembly. ACS Synth Biol 7(11):2665–2674.

48. Risso D, Ngai J, Speed TP, & Dudoit S (2014) Normalization of RNA-seq data using factor analysis of control genes or samples. Nat Biotechnol 32(9):896–902.

49. Bencivenga-Barry NA, Lim B, Herrera CM, Trent MS, & Goodman AL (2020) Genetic Manipulation of Wild Human Gut Bacteroides. J Bacteriol 202(3).

50. Altschul SF, et al. (1997) Gapped BLAST and PSI-BLAST: a new generation of protein database search programs. Nucleic Acids Res 25(17):3389–3402.

51. Katoh K & Standley DM (2013) MAFFT multiple sequence alignment software version 7: improvements in performance and usability. Mol Biol Evol 30(4):772–780.

52. Gilchrist CLM & Chooi YH (2021) clinker & clustermap.js: automatic generation of gene cluster comparison figures. Bioinformatics 37(16):2473–2475.

53. Sakamoto H, Kimura N, & Shimura Y (1983) Processing of transcription products of the gene encoding the RNA component of RNase P. Proc Natl Acad Sci U S A 80(20):6187–6191.

54. Kim S, Kim H, Park I, & Lee Y (1996) Mutational analysis of RNA structures and sequences postulated to affect 3’ processing of M1 RNA, the RNA component of Escherichia coli RNase P. J Biol Chem 271(32):19330–19337.

